# Single-cell perturbation dissects transcription factor control of progression speed and trajectory choice in early T-cell development

**DOI:** 10.1101/2021.09.03.458944

**Authors:** Wen Zhou, Fan Gao, Maile Romero-Wolf, Suin Jo, Ellen V. Rothenberg

## Abstract

In early T-cell development, single cells dynamically shift expression of multiple transcription factors (TFs) during transition from multipotentiality to T-lineage commitment, but the functional roles of many TFs have been obscure. Here, synchronized *in vitro* differentiation systems, scRNA-seq with batch indexing, and controlled gene-disruption strategies have unraveled single-cell impacts of perturbing individual TFs at two stages in early T-cell development. Single-cell CRISPR perturbation revealed that early-acting TFs Bcl11a, Erg, Spi1 (PU.1), Gata3, and Tcf7 (TCF1) each play individualized roles promoting or retarding T-lineage progression and suppressing alternative trajectories, collectively determining population dynamics and path topologies. Later, during T-lineage commitment, cells prevented from expressing TF Bcl11b ‘realized’ this abnormality not with a developmental block, but by shifting into a divergent path via bZIP and Sox TF activation as well as E protein antagonism, finally exiting the T-lineage trajectory. These TFs thus exert a network of impacts to control progression kinetics, trajectories, and differentiation outcomes of early pro-T cells.

## Introduction

Multipotent precursors establish T-cell lineage identity through a series of signal-modulated gene network steps(*1–5*), driven primarily by Notch ligands on the thymic stroma(*6–9*). Cells become committed to a T-lineage fate as pro-T cells, before T-cell receptor expression but multiple cell cycles after thymic entry(*4*). This intrathymic process can be mimicked in cell culture systems, e.g. OP9-DLL1 monolayer co-culture(*6*) and artificial thymic organoids, M-ATO (with DLL4)(*10*). Single-cell analyses have provided details of pro-T cells’ developmental trajectories(*11–13*); however, individual T cell precursors in the same intrathymic cohort can differ in developmental potentials, and timing is not deterministic in the T-lineage pathway(*14*). Multiple transcription factors (TFs) have been implicated in these events (rev. in (*15*)), but important questions remain how developmental kinetics and population distributions are regulated by these specific TFs within the early T-lineage gene regulatory network.

*In vitro* cell culture systems enable controlled timing of signaling environment encounters, crucial for interpreting experimental regulatory perturbations. However, during differentiation, cells still spread across heterogeneous developmental states. Therefore, we have combined specific TF perturbation strategies with scRNA-seq, using Cell Hashing for sample and batch indexing(*16*), to unravel the impacts of single TFs on cells at specific stages. We optimized a strategy based on a CRISPR/Cas9 knockout ‘perturb-seq’ system(*17*), exploiting the 10X Chromium V3 chemistry’s direct guide RNA (gRNA) capture capability(*18*), to resolve the detail of regulatory networks in primary ex-vivo derived early pro-T cells.

We have focused on two phases of pro-T cell programming (Fig. 1A): (1) the early gene regulatory network that guides hematopoietic precursors to initiate the T-cell program, and (2) the choices the cells undergo during commitment, under the control of the TF Bcl11b.

**Fig. 1.**
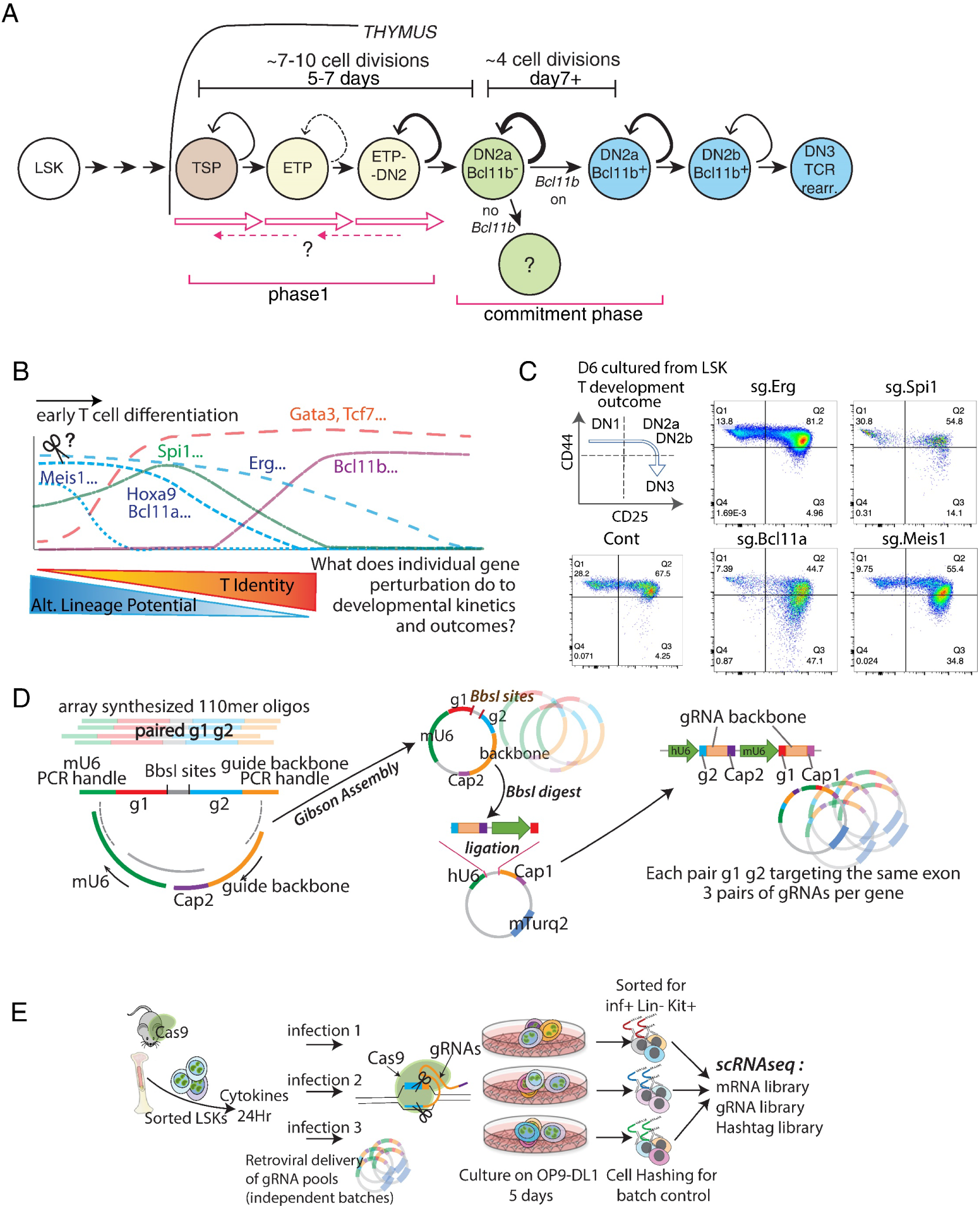
Developmental framework: Pool-based, batch-controlled TF perturbation scRNA-seq to define impacts of timed deletions of developmentally regulated TFs. (*A)* Experimental design and setup of internally controlled scRNA-seq experiments comparing wildtype (WT) and Bcl11b knockout (11b KO). (*B)* Gene expression dynamics of selected TFs in relation to alternative lineage potentials (*11*). (*C)* Representative surface expression phenotypes analyzed by flow cytometry. Acute TF deletions were induced in precursors before Notch signaling encounter (see Methods), then cells were cultured for 6 days with OP9-DLL1, and developmentally staged by surface expression of CD44 and CD25. Note: CD44 separates DN2a+b from DN3; DN2a is separated from DN2b by Kit expression(*2*) (not shown). (*D)* Pool-based dual gRNA cloning strategy used in the following experiments in this study. (*E)* Internal- and batch-controlled experimental setup for single-cell pool-perturbation RNA-seq. Two independently packaged stocks of the pool-perturbation viral library were each infected into separate cultures at MOI=1.0 and MOI=0.5, in parallel (plus an extra separate culture for one of the MOI=0.5 samples). The five resulting transduced cultures were each indexed with different hashtags before pooling for scRNA-seq analysis.

In early murine pro-T cells (Fig. 1A, phase 1), many multilineage and stem regulatory TFs are expressed overlapping with T-lineage TFs before lineage commitment(*11, 19, 20*). Some appear to retard T-lineage progression(*21–23*), and their T-cell program roles have been unclear. In the same cells, two important T-lineage regulators, TCF1 (encoded by *Tcf7*, referred to as “Tcf7” below) and Gata3 (rev. in (*5*)), are upregulated early, but their required viability roles(*24–28*) have obscured their precise impacts on the earliest lineage decisions.

Later, the TF Bcl11b is crucial for generation of αβ T cells and most γδ T cells, and its normal expression initiates precisely during commitment to the T cell lineage(*29–32*) (Fig. 1A, commitment). In pro-T cells lacking Bcl11b, bulk-population gene expression has shown numerous abnormalities. Populations retain some “immature” and lineage-inappropriate features while increasing diversion toward natural killer (NK)-like cells(*33–37*). However, the steps through which actual alternative pathways emerge have been obscure.

Here, scRNA-seq has distinguished perturbation effects on developmental progression speed from effects on developmental trajectories chosen in each of these contexts. In phase 1, a central kinetic opposition emerged between T-lineage-promoting and stem/progenitor-state preserving factors along a common trajectory, but distinctive roles for each TF were revealed. In contrast, Bcl11b-deprived cells around commitment did not regress backwards in differentiation trajectory but deviated to an alternative, divergent developmental trajectory. This approach clearly distinguishes TF impacts on developmental speed from TF effects on trajectory choice. In addition, SCENIC regulon analyses reveal novel, latent TF participants at developmental branchpoints that become activated in specific mutants.

## Results

### Targeting of TFs with early dynamic expression patterns through batch controlled, dual-gRNA direct-capture perturb-seq

To reveal key controllers of gene expression changes before T lineage commitment, we examined the effects on differentiation speed and outcomes when we knocked out (KO) any of seven candidate regulators of these events (Fig. 1B). To control this process and the timing of the knockout, we used two in vitro differentiation systems, OP9-DLL1 coculture(*6*) and ATO-mDLL4 cultures(*10*). We first carried out extensive control experiments to validate their use, comparing them with intrathymic (Thy) cells(*11*) in terms of differentiation timing and transcriptome changes (Methods) (Fig.S1A). Both bulk RNA-seq of DN1, DN2a uncommitted, and DN2a newly committed populations (sub-divided by Bcl11b-YFP expression onset(*32*))(Fig. S1; Table S1) and scRNA-seq analyses (Fig. S2A-G) supported the comparability of in vitro and in vivo intrathymic differentiation.

Based on preliminary studies suggesting effects on differentiation, we focused on *Bcl11a, Spi1, Hoxa9, Meis1*, and *Erg,* which encode stem or progenitor-associated factors, and on *Tcf7* and *Gata3*, which encode T lineage-associated factors (Fig. 1B). KO effects were elicited by using retroviral vectors to introduce single guide RNAs (sgRNAs) against the gene of interest into ROSA26-Cas9-transgenic LSK cells, purified from mouse bone marrow (BM). Although several targeted genes are important for viability of early T cell precursors(*25, 28, 38–40*), we ensured maximum survival of perturbed cells by using *Bcl2-tg;ROSA26-Cas9* input cells. After transduction, the cells were then transferred to culture to initiate the T cell program.

In preliminary studies, KOs of progenitor-related genes, *Bcl11a* or *Meis1*, suggested an accelerated DN2a to DN2b transition, while *Erg* KO apparently increased the rate of DN1 to DN2 transition, as indicated by surface markers CD44, c-Kit and CD25 (e.g., Fig. 1C). Previously, disruption of *Spi1*, encoding PU.1, a pioneer factor important for the myeloid vs lymphoid decision, was found to accelerate DN2 to DN3 progression while reducing cell yield(*38, 39*). Here, even with the *Bcl2* transgene to improve recovery, earlier deletion suggested similar effects. However, inter-culture variation was considerable. Our highly-controlled scRNA-seq strategy (Fig. 1D-E) was designed to combat variability and non-autonomous effects in determining whole-transcriptome effects of these perturbations, using a pool-synthesized, batch-controlled dual gRNA system.

Briefly, Cas9^+^ Bcl2^+^ precursor cells (purified LSK) were infected at low multiplicity with a library of vectors, each including paired gRNAs against the same target gene. The cells were then cultured on OP9-DL1 stroma for 5 days before sorting for infection-positive populations, which were then analyzed through scRNA-seq (Fig. 1E; Methods). The experimental conditions of 5-day-cultures of LSK with Notch signaling were chosen to monitor loss of function effects in phase 1, before T lineage commitment. The paired gRNAs (each pair designed against the same exon of the same gene to ensure sufficient KO) were synthesized through array-based oligo synthesis, then the oligo pool was PCR amplified and Gibson-assembled to generate the paired dual gRNA insert pool, and subsequently incorporated into retroviral backbones and packaged into the final retroviral vector library (Fig. 1D). The paired gRNAs being expressed were compatible with direct capture of both gRNA sequences in scRNA-seq using 10X Chromium V3 chemistry (‘Cap1’ and ‘Cap2’ in Fig. 1D; for details, quality controls, validation tests, and titration, see Methods; Figs. S3A-D).

Importantly, the whole library was prepared in two independent packagings which were each separately infected into BM cells and incubated in parallel, separate replicate cultures (Fig. 1E; see Methods). After 5 d incubation, these replicates were harvested and each labeled with distinct antibody-conjugated “cell-hashing” oligonucleotides(*16*), and then pooled for scRNA-seq analysis. After sequencing libraries of the cDNA, gRNA, and hashtags from the same experiment, the resulting FASTQ files were aligned using CellRanger3 (for cDNA) and an in-house pipeline (Fig S3E,F; Methods). The multiple replicate infections, multiple sgRNA pairs against the same target, and pooled sequencing of five separately cultured replicates in the same 10X v3 analysis yielded a high-quality, internally-controlled data resource in which to identify specific perturbation effects.

### Dramatic changes in topology with single TF perturbations

In scRNA-seq data, the earliest and latest phase 1 cells in a low dimensional transcriptome representation (Fig. 2A-C) were identified by the expression of previously determined landmark genes(*11*) (Fig. 2B; Fig. S4A; top cluster markers listed in Table S2). Some individual TF perturbations resulted in notable pattern changes (Fig. 2C,D). While control cells spread sparsely across the UMAP1 and 2 space (Fig. 2D left), cells in the same population expressing sg.Tcf7 stalled at the most un-differentiated stage, whereas cells expressing sg.Spi1 shifted towards more differentiated stages, slightly veering from the control trajectory (Fig. 2D). Effects of sg.Gata3, sg.Bcl11a, sg.Hoxa9 or sg.Meis1 were subtler (Fig. 2E). But cells expressing sg.Erg formed a distinct cluster parallel to the normal trajectory (Fig. 2D). Despite matched inputs, more sg.Erg expressing cells (all sg.RNA pairs) emerged in the pool than any other cells. This suggested that Erg loss may enhance proliferation (Fig. 2F). In contrast, Tcf7 and Meis1 perturbations significantly reduced cell yields despite the anti-apoptotic Bcl2 transgene (Fig. 2F). Most noted effects were seen with at least 2/3 of the sg.RNA pairs for a given gene, although there were some construct-specific differences (Fig. S4B). Conservatively, below, we take the averaged effect of all sg.RNAs against the same gene as a minimal estimate of the ‘KO’ impact.

**Fig. 2.**
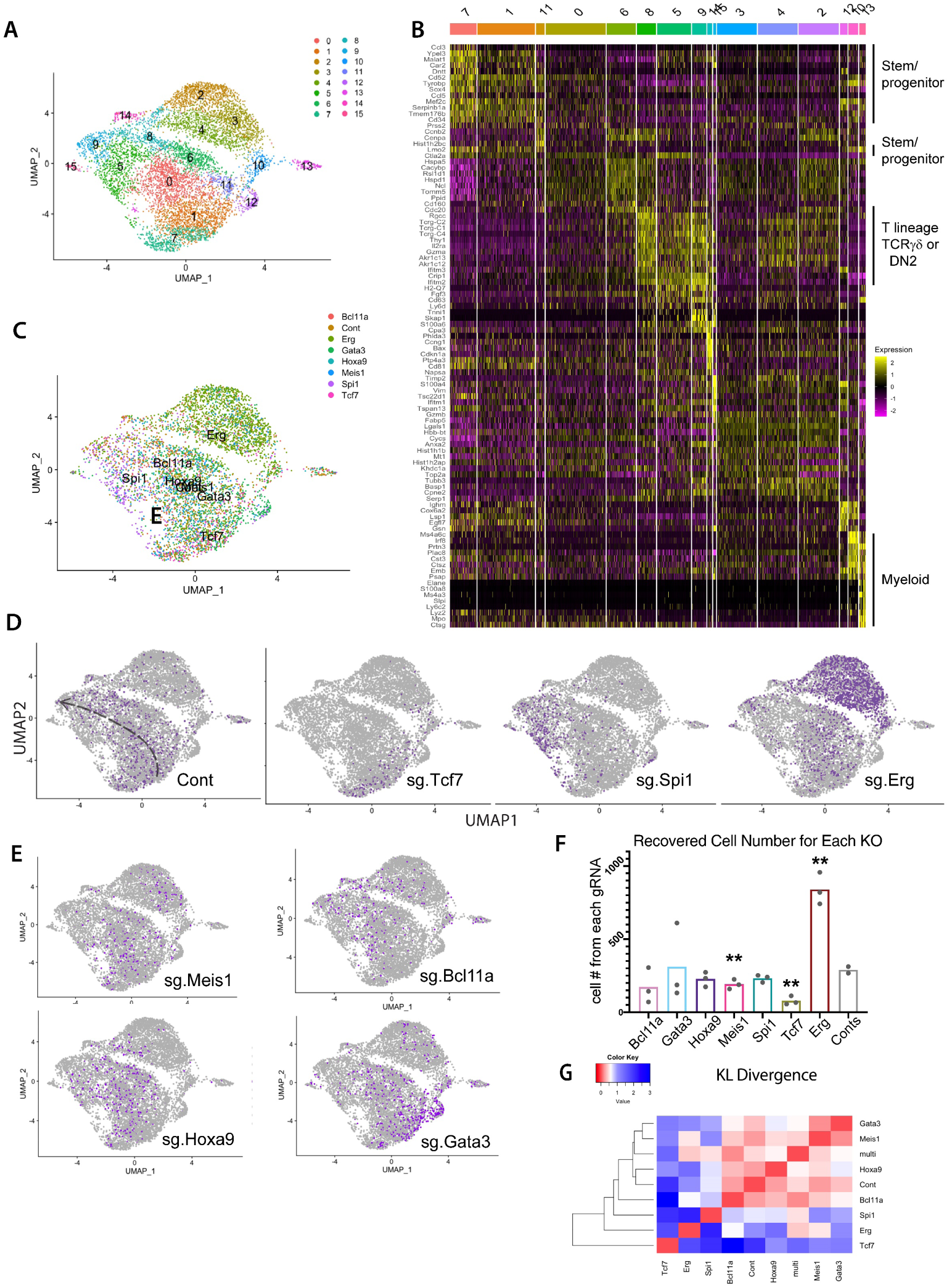
Impacts of TF perturbations on gene expression, cell number, and differentiation trajectories of early pro-T cells. *(A)* UMAP 1-2 display of scRNA-seq data from total set of pool-perturbation samples, here based on PC 1-16. N= 5541 cells for singly assigned gRNA vector, N=8045 for total recovered single cells. The cells are colored by Louvain clustering algorithm. (*B)* Heatmap displaying the top 10 enriched genes in each sub-cluster, in approximate developmental order. Defined by Seurat 3 pipeline with minimum fraction of expressing cells ≥ 0.25, Wilcoxon rank sum test with avg_logFC threshold of 0.3, full list of genes in Table S2). (*C-E)* Cells in UMAP display, colored by sgRNA assignment. All cells with any of the 3 pairs of dual sgRNAs against the same gene are colored together. Number of cells of each KO: Controls (Cont.): 793, Bcl11a: 353, Erg: 2101, Gata3: 686, Hoxa9: 503, Meis1: 409, Spi1: 481, Tcf7: 215. (*C)* Merged representation of results with labels showing the centroids of different KO distributions on UMAP1-2. (*D-E)* Cells from the indicated KOs (purple dots) within the total sample (gray dots). Trajectory was manually sketched based on marker gene expression shown in panel *(B)*. (*F)* Cell numbers recovered from the scRNA-seq pool. Each dot represents the cell number recovered from one of paired gRNA vectors against the indicated target. Statistical significance calculated in individual pairwise t-tests between Control and each of the KOs. (**: p-val<0.01, *: p-val<0.05). (*G)* Heatmap of KL divergences (red: high similarity; blue: low similarity) among all sample types, WT and aggregated KO for each gene.

Pair-wise Kullback-Leibler (KL) divergences among the cluster distributions of each genotype showed that KOs of *Tcf7*, *Erg*, and *Spi1* each gave dramatically distinct cluster distribution patterns, very different from controls (Fig. 2G). However, except for the clusters specific for the Erg KO cells, most of the KO and control cells were still found within the same ‘common’ transcriptome clusters, although their distributions among these clusters shifted greatly (Fig. 2C; Fig S4A). Within the same shared clusters, cells from different genotypes also had closely matching gene expression patterns (Fig. S4C,D). In contrast, the three Erg KO-specific clusters (clusters 2-4, Fig. 2A, C, D; Fig. S4A) showed a distinct gene expression signature unlike the ‘early’ clusters (1, 7, 11) or the ‘late clusters (5, 6, 9)(Fig. 2B).

### Shifts in T-cell differentiation pseudotime trajectory caused by KO of individual TFs

The impacts of these TF KOs on developmental progression could be related to a common T-lineage differentiation pseudotime metric. Trajectories were calculated from full transcriptome profiles (Monocle3 analysis) in 3D UMAP space (Fig. 3A, pseudotime progressing from dark color to light color). The single dimension of pseudotime indicated distinctive KO impacts on gross developmental “speed” (Fig. 3B; red bars, mean). Bcl11a, Spi1 and Erg KOs showed significant pseudotime acceleration relative to control (Kruskal-Wallis test; significance asterisks in blue), while the Tcf7 KOs gave significant deceleration from control (significance asterisks in red). Gata3 KO cells progressed beyond Tcf7 KO cells, but specifically failed to generate cells matching the most advanced controls; Hoxa9 and Meis1 KO effects were less significant. Thus, the natural expression of *Spi1*, *Bcl11a*, and *Erg* in early pro-T cells appears to retard the differentiation process actively, while expression of *Tcf7* and *Gata3* is used (or required) to advance the T lineage program, consistent with previous studies(*24, 28, 39*).

**Fig. 3.**
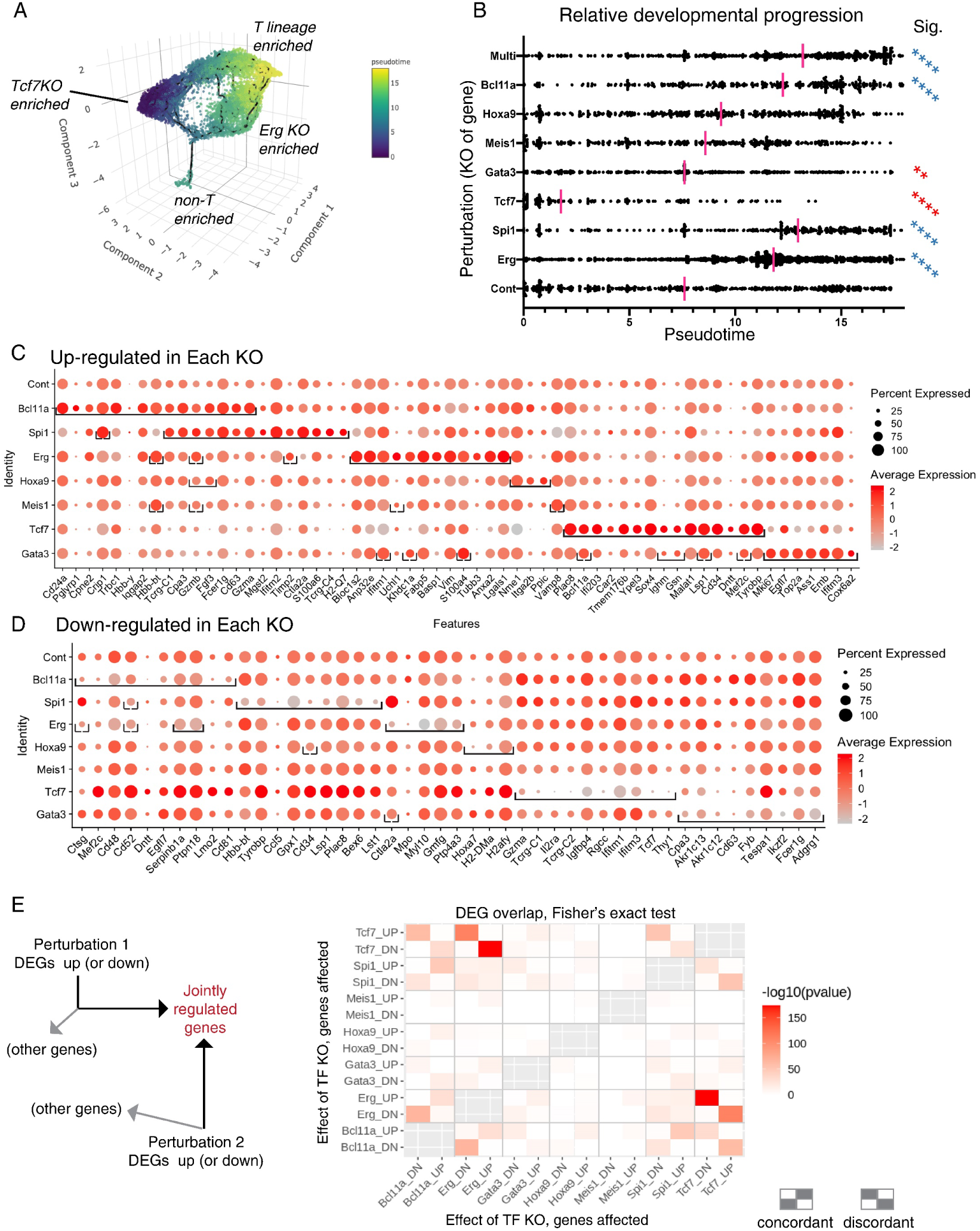
Impacts of TF knockouts on gene regulation, pseudotemporal progression, and discrete regulon modules in early pro-T cells. *(A)* 3D UMAP of all samples, colored by inferred pseudotime, to relate cells to a common overall pseudotemporal advancement score regardless of trajectories. The pseudotime was calculated with Monocle 3, based on the trajectory inference from 3D UMAP built using the size and cell cycle scaled data as described in Fig. 2A (see Methods). (*B)* Pseudotime distributions (x axis) of cells from the indicated KOs (category axis), with means indicated (red bars). The small “non-T-enriched” population seen in *(A)* was excluded from the calculation. Statistical significance of comparisons, each KO to control, by Kruskal-Wallis test of multiple comparisons. **, adj.p-val < 1E-02; ****, adj.p-val < 1E-04. Blue asterisks: faster, red asterisks: slower than Control (Cont). (*C-D)* Genes highly differentially expressed between control and KOs, full gene lists in Table S3. (*C)* Top 15 up-regulated genes (x axis) for each KO (y axis). Top 15 for Bcl11a KO are followed by top 15 for Spi1 KO if not already listed, etc. Brackets enclose the top differentially expressed genes for that KO; some bracketed genes are listed before others if they are also upregulated in another KO above. Top 15 defined by avg_logFC function on Seurat 3-processed data. Significance determined by Wilcoxon Rank Sum test, minimum expression fraction ≥ 5% cells in either the control or the KO, average “log expression” difference ≥ 0.1 (ln, baseline pseudocount=1), and adjusted p-val < 1E-02. (*D)* Top 10 down-regulated genes in each KO, as in panel (*C*), with average “log expression” difference ≤-0.1, and adjusted p-val < 1E-04. *(E)* Pairwise comparisons of top differentially expressed genes in each condition, by Fisher’s exact test.

### Factor-specific effects of different KOs at target gene level

All KOs except Hoxa9 and Meis1 significantly changed expression of many genes (top hits shown as dots within brackets in the row for that KO in Fig. 3C, D; extended data in Table S3). The Bcl11a KO down-regulated progenitor-associated genes (e.g. *Lmo2, Mef2c, Egfl7*)(*11*) and modestly upregulated *Gata3* and *Tcf7,* and also NK and ILC-associated genes (*Zfp105, Zbtb16, Fcer1g, Gzma, Gzmb*). The Spi1 KO overlapped with the Bcl11a KO’s differential target expression, and included increased expression of T-cell receptor-associated genes (*Tcrg-C1, Tcrg-C4, Trdc, Trat1, Cd247*). Erg KO cells also downregulated progenitor genes, but upregulated distinctive cytoskeleton- and growth/signaling-related genes. Tcf7 KO cells lost expression of T-cell genes but showed expression enriched for stem/progenitor-cell related genes(*11*) (*Sox4, Cd34, Lmo2, Bcl11a, Spi1*, *Mef2c*) and lncRNA *Malat1*. While Gata3 KO cells also enriched expression of some progenitor genes, they showed a different pattern from the Tcf7 KO; instead, they upregulated a mosaic of genes enriched in different non-T cell types, as well as cell cycle-related genes.

We related the top genes affected by each KO quantitatively to the ‘normal’ maturation-related gene-expression shift in the control samples between early DN1 to late DN2b, i.e., genes changing expression between early (7,1,11) and late clusters (0,6,5,8,9,14) in Control samples only (Fig. 2B; Fig. S4E). Volcano plots (Fig. S4F-I) show that the Tcf7 KO effect indeed resembled a simple developmental block (Fig. S4F) relative to genes normally up-regulated (magenta) or down-regulated (cyan) in T-lineage development. While Spi1 or Bcl11a KOs caused effects roughly opposite to Tcf7 KO, Gata3 KOs affected maturation-related genes irregularly, and both Gata3 and Spi1 KOs strongly affected genes that could not be related purely to the normal T-lineage trajectory (Fig.S4G-I, dark brown dots). Thus, these regulatory effects were TF-specific, not simply consequences of developmental progression states.

Pairwise comparisons showed that when Spi1 KO and Bcl11a KO affected the same genes, they acted concordantly nearly all the time, whereas when Tcf7 KO affected the same genes as Bcl11a KO or Spi1 KO, it nearly always affected them in the opposite direction, as shown in a heat map of Fisher’s exact test comparisons in Fig. 3E. This was consistent with *Tcf7* acting normally to promote T-lineage progression while *Spi1* and *Bcl11a* normally opposed progression. However, Gata3 KO effects were mixed, and Erg KO effects cut across this developmental distinction. Its effects were predominantly antagonistic to Tcf7 KO, but more concordant with Bcl11a KO than with Spi1 KO (Fig. 3E). Thus, despite similarities in the one-dimensional pseudotime patterns of response to different KOs, in fact each TF exerted its own distinctive effects on T lineage progression.

### SCENIC regulon analysis: candidate indirect mechanisms contributing to developmental programs

While some effects could be direct, these TFs might also control some targets indirectly, as multiple KOs affected expression of other regulators. Seurat 3-defined DEGs indicated mutual repression between *Bcl11a* and *Gata3*, positive regulation of *Mef2c* and *Lmo2* by Erg, and opposing regulation of *Bcl11a*, *Mef2c*, and *Lmo2* by Spi1 and Tcf7 (Table S3). To gain broader insight into the most influential intermediate regulators, we used the SCENIC algorithm(*41*) to identify groups of genes potentially co-regulated directly by activities downstream of the seven TFs. SCENIC infers candidate direct targets independent of perturbation, based on enrichment of a well-defined TF target motif in genomic regions near the promoters of genes that co-vary with expression of that TF. Such potential positively-regulated target gene sets are defined as ‘regulons’. Not all TFs of interest had well-defined regulons, but SCENIC identified many regulons with sharply varying activity patterns across the dataset (Fig. 4A; Fig. S5A-F). These generally agreed with the expression of the corresponding TFs from ETP-DN2b stages (Fig. 4A; cf. (*11, 42*)). Notably, a regulon’s enrichment sometimes appeared more restricted than its corresponding TF. For example, Nfkb1 regulon activity (Fig. 4A, center) was enriched in later DN2-stage cells, although most NF-κB/Rel family members are expressed from early stages (Fig. S5G). Thus, SCENIC might also indicate when signaling pathways controlling factor activity post-transcriptionally are an additional rate-limiting constraint.

**Fig. 4.**
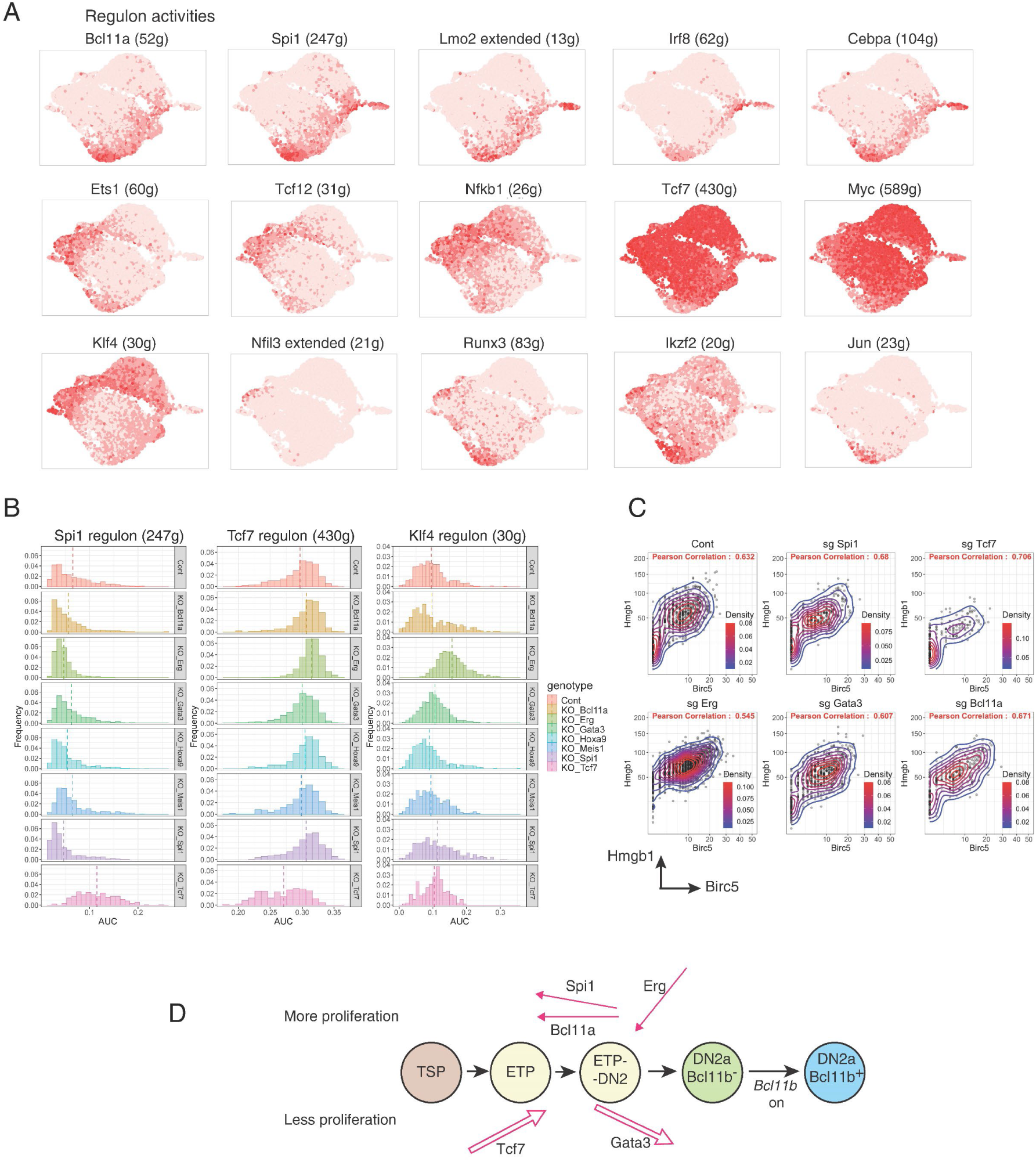
SCENIC analysis of major regulons in early pro-T cell pool-perturbation dataset and impacts of TF knockouts. *(A)* Activities of inferred regulons for different endogenous TFs across the whole pool-perturb early pro-T cell dataset. Color intensity in UMAP indicates expression of genes constituting the regulons predicted to be activated by the indicated TFs (gene lists in Table S4B). Top: progenitor-associated. Middle: associated with normal T-lineage progression. Bottom: not normally associated with T-lineage progression. *(B)* Histograms of activities of the Spi1, Tcf7, and Klf4 regulons among cells from the 8 perturbation conditions (“genotypes”). Plots show frequency (y axis) vs. area under the curve (x axis) for each regulon within the indicated subset. (*C)* Pairwise scatterplots of the *Hmgb1* and *Birc5* cell cycle-associated gene transcripts among the different KOs (G2/M phase cells express *Birc5*). (*D)* Summary schematic of inferred physiological roles of Tcf7, Gata3, Spi1, Bcl11a, and Erg in developmental progression (left to right) and proliferation control (up vs. down) in normal early pro-T cell development.

As expected, regulons for Spi1 and Tcf7 TFs showed specifically reduced activity in the respective KOs (Fig. 4B, Table S4; stars in Fig. S5A). In addition, in accord with a global developmental progression block, Tcf7 KO cells were enriched for expression of all progenitor-associated regulons, including the Spi1, Bcl11a, Irf8 and Cebpa regulons (Fig. 4B, Fig. S5A, E), Table S4). In contrast, Spi1 KO increased activity of the predicted Tcf12, Ets1, and Nfkb1 regulons, associated with T-lineage progression, suggesting that PU.1 encoded by *Spi1* normally holds these TF activities in check (Fig. S5C, Table S4). Most notably, the Erg KO caused highly specific upregulation of the Rxra and Klf4 regulons (Fig. 4A, bottom; Fig. 4B; Fig. S5B, D; Table S4), although Klf4 is normally restricted to ILC and myeloid cells(*42*). Thus, Erg could normally prevent expression of Klf4 or a related factor. In summary, SCENIC gave substantial evidence for cascading regulatory network impacts.

### Cell cycle and RNA content control distinct from developmental progression

RNA contents, S phase and G2/M scores differed among the various KOs (Fig. S6A-C), with the Erg KO and Tcf7 KO at opposite extremes. SCENIC identified a Ybx1 regulon prominently associated with cell cycle in all samples, including mitotic spindle genes and S/G2 cell cycle markers, e.g. *Birc5* and *Hmgb1*. The Ybx1 regulon was also predicted to include *Myc*, encoding a TF with potent metabolic effects and its own large regulon (Fig. S5F). Despite their opposite effects on development, the Spi1 and Tcf7 KOs both reduced cycling indices and reduced expression of the Myc regulon (Fig. S5F)(cf. refs.(*39, 43*)). In contrast, the Erg KO shifted cells to a more cycling state, with high Ybx1 activity and slightly increased activity of the Myc regulon (Fig. 4C, Figs. S5F, S6D), consistent with the high cell yield. Interestingly, Gata3 KO cells resembled Erg KO more than Tcf7 KO cells in upregulation of cell-cycle genes *Mki67, Top2a, Birc5* and *Hmgb1* (Figs. 3C, 4C; Tables S3, S4).

SCENIC analysis thus confirmed that perturbation effects included TF-specific modular responses beyond the simple dichotomy between developmental progression-promoting and progression-opposing actions. Highly TF-specific effects on cell cycle cut across this dichotomy, and different KOs unleashed activities of different alternative pathway regulons. The asynchronous, stochastically progressing differentiation of the early pro-T cells emerged from these distinct forces (Fig. 4D).

### Routes and kinetics of lineage divergence of Bcl11b KO pro-T cells

The culmination of early pro-T cell development is lineage commitment, occurring as *Bcl11b* is finally activated(*11, 32*) (Fig. 1A, Fig. S4A). From its first expression, Bcl11b is required to preserve lineage fidelity(*30, 37, 44, 45*); however, the aberrant cells that emerge when Bcl11b is deleted have obscure origins. Previous bulk studies(*34, 37*) have shown that Bcl11b KO pro-T cells not only express some early ‘immature’ or ‘non-T’ associated genes normally downregulated during commitment, but also abnormally turn on genes that were silent in normal precursors before commitment, i.e. before Bcl11b was there to repress them. We therefore investigated the trajectories and regulatory components through which Bcl11b loss causes cells to activate such alternative differentiation pathway(s).

We exploited the longer-term differentiation fidelity of the ATO system (Fig. S1) to compare WT and Bcl11b KO precursor differentiation at single-cell resolution. Because *Bcl11b*^-/-^ pro-T cells have no survival defect in vitro(*28, 33*), for the KO we simply used conditional knockout *Bcl11b^flx/flx^*;*Vav1-iCre* mice(*37*). These delete exon 4 in *Bcl11b* before T cell development begins; but deletion should only affect cells after ∼D7 in the ATO system when *Bcl11b* expression is normally activated. Multiple parallel ATOs were seeded with LSK precursors purified from different individual wildtype (WT) or Bcl11b KO mice, and at D10 and 13 the Lin^-^ CD45^+^ CD25^+^ cells were sorted, separately, from different individual mice as replicates (Fig. 5A). The surface staining phenotypes consistently recapitulated phenotypes of Bcl11b KO pro-T cells from the thymus, with Bcl11b KO cells exhibiting higher c-Kit surface staining than WT (Figs. 5B, S7A)(*33–37*). Cells derived from different individual animals were cultured separately and each tagged with cell hashing antibody conjugates to associate different barcodes with each donor(*16*), then pooled for scRNA-seq within a single 10X v3 run (see Methods). The results showed excellent reproducibility across samples and in two separate experiments (Fig. S7B-D).

**Fig. 5.**
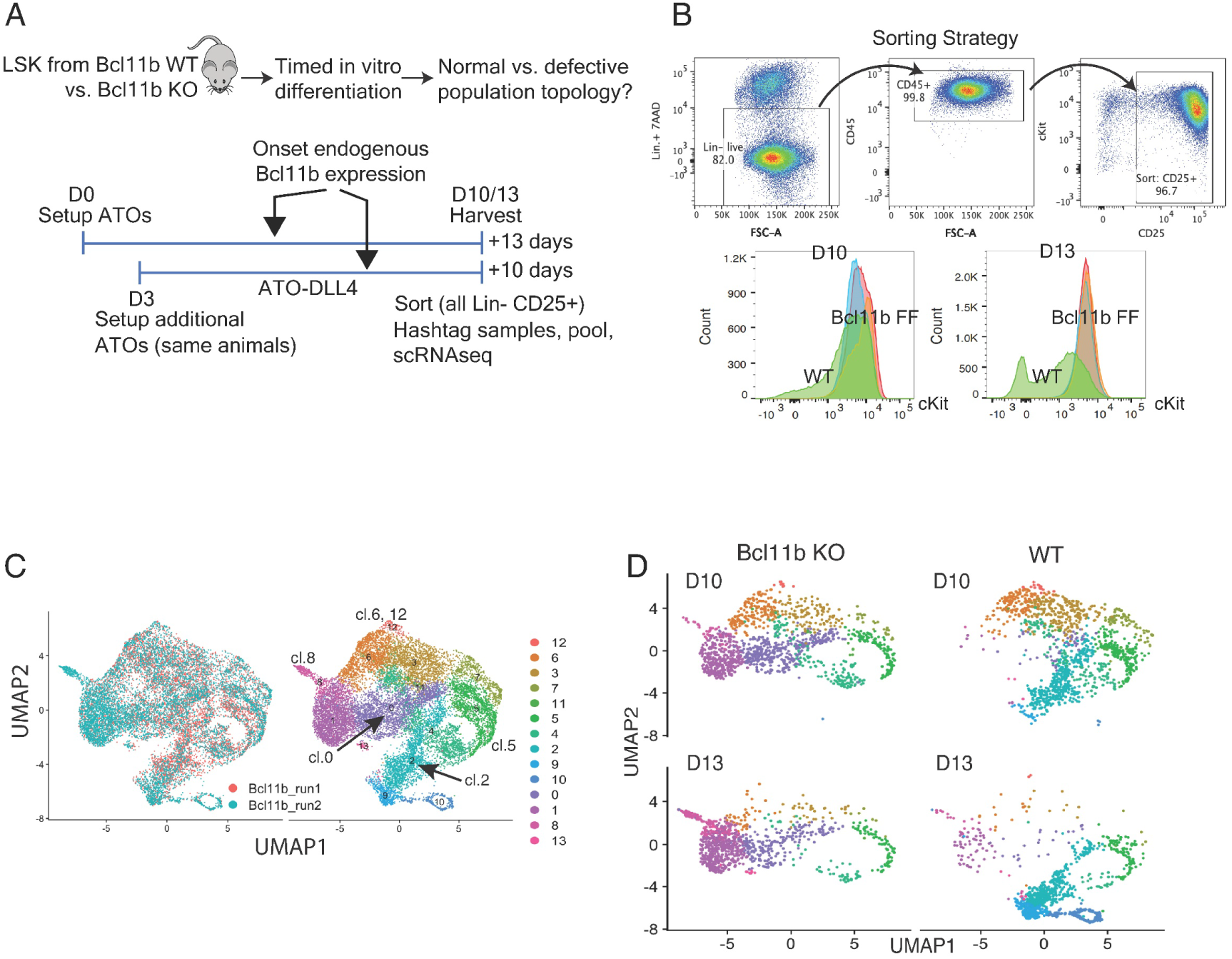
Time-dependent progressive divergence of Bcl11b knockout from wild type gene expression trajectories at late pro-T cell stages. *(A)* Schematic of experiments to determine pathway of response to loss of Bcl11b. Arrows indicate that endogenous *Bcl11b* normally turns on at approximately D7 of differentiation. (*B)* Top: FACS purification strategy for the Bcl11b scRNA-seq experiments. Bottom: surface staining phenotypes of progenitor-marker c-Kit levels in WT (green) and Bcl11b KOs (replicates in blue, red, orange) (noted as FF for flx/flx, details in Methods). (*C)* Left: alignment of two experiments of Bcl11b KO scRNA-seq profiles after CCA scaling, as ‘Bcl11b_run1’ and ‘Bcl11b_run2’, in UMAP1-2. N=7451 cells from ‘Bcl11b_run1’, and N=8558 cells from ‘Bcl11b_run2’, total 16009 cells. Note that ‘Bcl11b_run1’ samples were only collected in D10. Right: Louvain clustering of the integrated samples (see Methods). *(D)* Samples subsetted according to the hashtag demultiplexed genotype and time of harvest, and displayed in UMAP1-2, colored by the same clustering annotation as in panel F (individual mouse samples shown in Fig. S7B).

### Shifts in pattern in low-dimensional space and increasing divergence with time

Single-cell transcriptomes of Bcl11b KO and WT cells, displayed in UMAP 1-2 (Fig. 5C), showed partial overlap between Bcl11b KO and WT cell clustering patterns at D10, but gave strikingly different patterns at D13 (Fig. 5D, Fig. S7B). For these samples, the UMAP topology was most instructive without cell cycle regression; S+G2M-enriched cell clusters appear to the right. The main clusters (Fig. 5C) could be identified by their gene expression patterns (Fig. 6A, B, Table S5, cf. (*11*)). Appearance of *Bcl11b* locus transcripts was clearly visible as a developmental milestone in WT and Bcl11b exon-disrupted cells, alike (Fig. 6B). At D10, cultures of both genotypes still included many cells in immature states (clusters 12, 6, 3) as well as intermediate cells strongly expressing cell cycle genes (clusters 7, 5). By D13, the immature clusters were depleted in both genotypes (Fig. 5G). However, as WT cells increasingly shifted “down” to more mature (‘Comm. T’) stages in clusters 4, 2, 9, and 10, the Bcl11b KO cells shifted “left” to alternative ‘aberrant KO’ clusters 0, 1, and 8 instead (Fig. 5G; genes shown in Fig. 6A). Cluster 8 especially, the “10 o’clock” tip of the UMAP plots, expressed Bcl11b KO-specific genes with little if any normal expression in pro-T cells (Fig. 6B, middle and bottom rows), described below. Pair-wise KL divergence calculations between all the samples based on each sample’s cluster distributions (Fig. 6C) confirmed the pattern agreements between all the experimental and biological replicates and the marked contrast between WT and Bcl11b KO samples, with increasing divergence between WT and Bcl11b KO patterns with time.

**Fig. 6.**
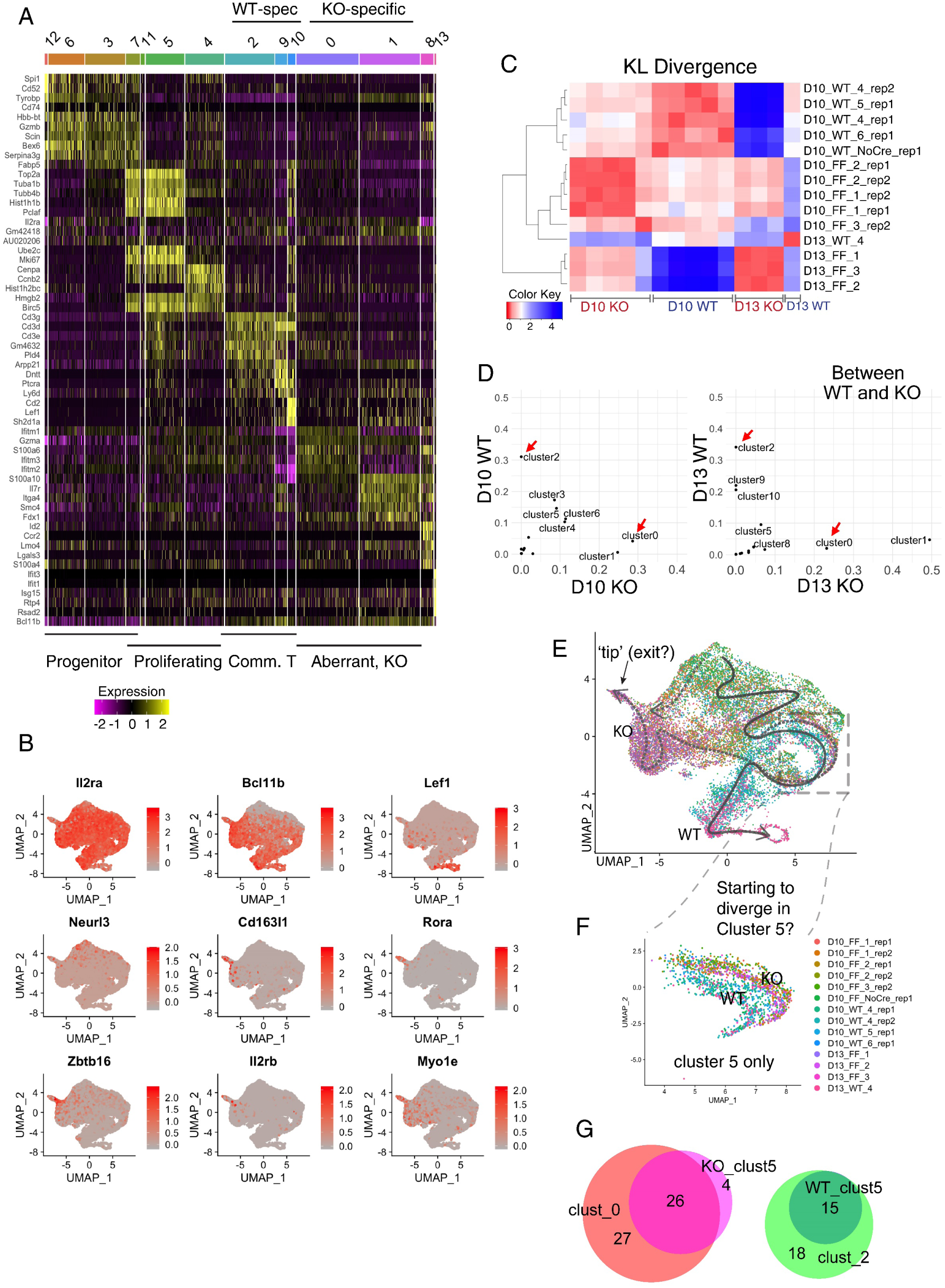
Single-cell analysis of differences between Bcl11b KO and WT: WT and Bcl11b KO trajectories separate immediately after the normal onset of Bcl11b expression. *(A)* Heatmap displaying the top 5 enriched genes in each sub-cluster ordered by approximate developmental trajectories of WT and Bcl11b KO, based on gene expression and connectivity in UMAP displays. (Seurat 3 pipeline, minimum fraction of expressing cells ≥ 0.25, Wilcoxon rank sum test, avg_logFC ≥ 0.3). *(B)* Expression patterns of indicative genes in UMAP 1-2 plots of whole WT and Bcl11b KO data ensemble. Top row: genes indicating degrees of progression in the normal pro-T cell pathway. Note that transcripts from *Bcl11b* locus are detected even from mutant allele. Middle and Bottom rows: genes distinctively upregulated in Bcl11b KO cells. *(C)* Heatmap showing the KL divergences among all of the integrated samples, calculated based on correlations between cluster distributions (Table S6B). *(D)* Pair-wise cluster distribution scatterplots comparing WT and Bcl11b KO. Pearson correlation r= −0.04 for D10, r= −0.18 for D13. Red arrows indicate the most dramatic and consistent differences between the two genotypes in both time points, cluster 0 and 2. *(E-F)* UMAP 1-2 colored by individual demultiplexed samples. *(E)* Schematic, overview of inferred trajectories of differentiation of WT and Bcl11b KO cells around stages of commitment. *(F)* Zoom-in view of subset of cells only within cluster 5 from Fig. 5F, showing the slight separation between genotypes. N=1630 cells. *(G)* Venn diagrams comparing gene expression changes between WT and Bcl11b KO within cluster 5 with those in definitively divergent clusters 0 and 2 (cf. panel D).

Genes changing expression from D10 to D13 clearly differed between Bcl11b KO and WT populations (Table S6; Fig. S8A, B). To identify the most quantitatively important transcriptome clusters that were diagnostic of the genotype differences, we used data from experiment 2, where the two timepoints were collected from each of the same inputs (see comparison, Fig.S7C-F). At both D10 and D13, cluster 2 was always enriched as a proportion of WT cells and cluster 0 was always enriched in the KO (Fig. 6D), thus indicating an early stage of WT and Bcl11b KO trajectory separation at D10. The top 20 genes differentially enriched between cluster 2 and cluster 0 are shown in Fig. S8C (full list in Table S7A). Cluster 2-enriched genes were mostly T lineage-associated, while the cluster 0 enriched genes were enriched for previously reported Bcl11b repression targets(*34, 37*). While some KO-specific genes were shared with the ‘immature’ clusters (12, 6, 3), a similar proportion appeared to be upregulated uniquely in the Bcl11b KO clusters, emphasizing a distinct program. These genes were clearly differentially expressed between clusters 0 and 2, and became even more differentially expressed further down the ‘branches’ (clusters 2, 9, 10 for WT and 0, 1, 8 for Bcl11b KO, respectively). These gene expression analyses suggested likely differentiation trajectories, shown schematically in Fig. 6E.

### Initiation of divergence in highly cycling cells around time of normal *Bcl11b* activation

Close comparison of the profiles in Fig. 5F,G suggested that cluster 5 could also harbor early Bcl11b-dependent differences (box in Fig. 6E). This cluster, roughly corresponding to when the *Bcl11b* locus is first transcribed, was represented in both genotypes at both timepoints (Fig. 5G). It was enriched for cell cycle-associated genes (Fig. 6A) and formed a loop-like topology (with cluster 4) suggesting a proliferative amplification phase, that was well defined in these plots of data without prior cell cycle regression. Interestingly, within cluster 5, the WT and Bcl11b KO cells formed parallel but slightly separate loops (Fig. 6F). Differential gene expression analysis between WT and Bcl11b KO cells within cluster 5 only (Fig. S8D, Table S7B) showed numerous genes already significantly differentially expressed within this seemingly mixed ‘amplification loop’, before the cells split overtly into cluster 0 and cluster 2. These genotype-differentially expressed genes within cluster 5 already represented nearly half of all genes that became differentially expressed between cluster 0 and cluster 2 (Fig. 6G). Moreover, even within cluster 5, the WT to KO difference increased between D10 and D13. For example, at D10, expression levels of *Ikzf2*, *S100a10,* and *Itga4* in Bcl11b KO cells in cluster 5 were similar to levels in the WT cells, but by D13 these genes were substantially upregulated in Bcl11b KO cells in cluster 5, compared to WT cells in cluster 5 (Fig. 7A, red dots). Thus, the initial distinction between WT and Bcl11b KO emerged during the shared proliferative phase represented by cluster 5, around the time when the *Bcl11b* locus is normally turned on (Fig. 6B). In such a lineage branchpoint, the accumulation of subtle abnormal expression features could precede substantial movement between discrete transcriptome clusters in low dimensional space.

**Fig. 7.**
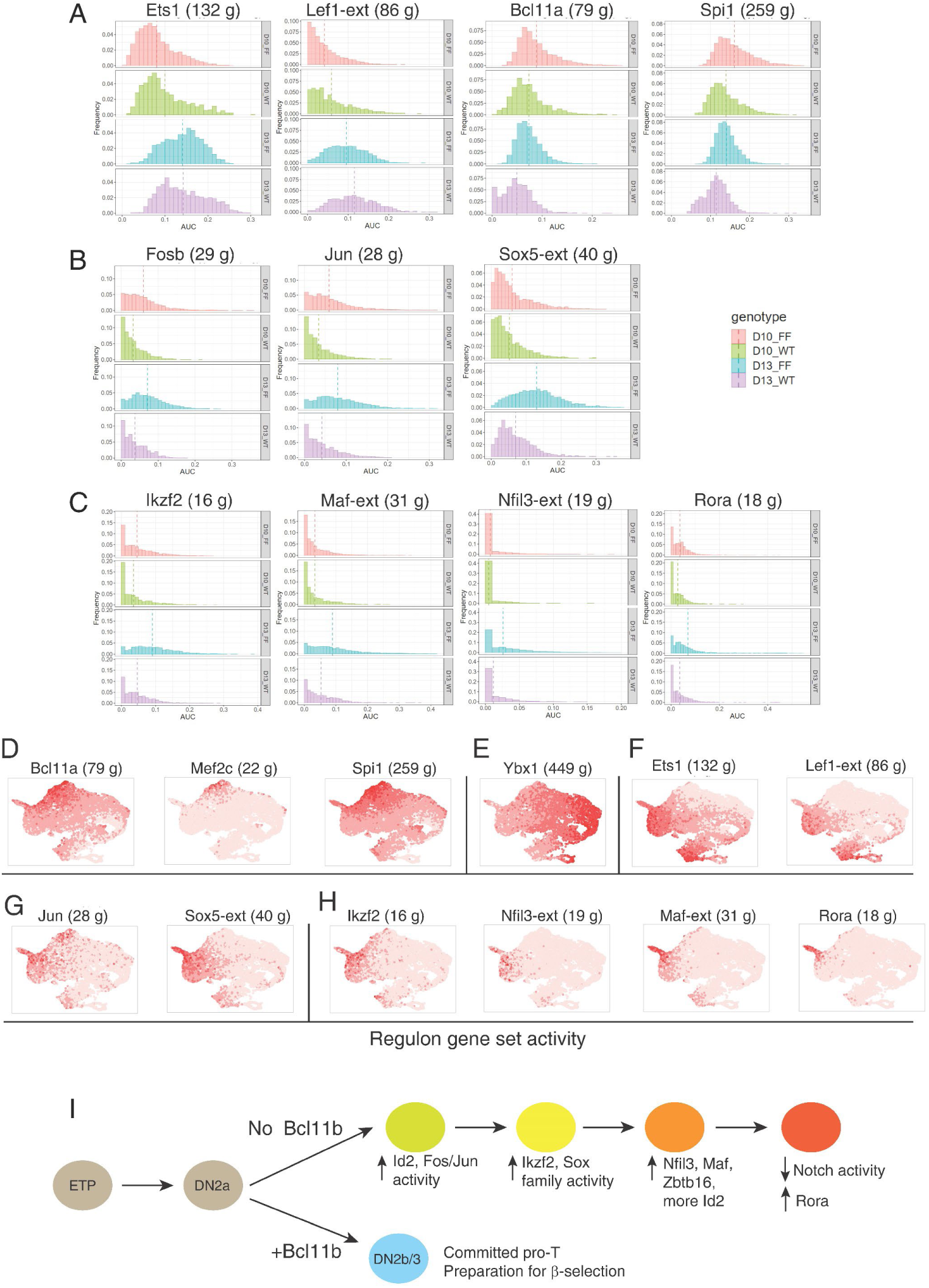
SCENIC analysis of Bcl11b KO impact on gene expression: potential intermediates from AP-1 family implicated in an alternative gene network cascade. *(A)-(C)* Histograms of expression enrichments for indicated regulons, comparing cells from Bcl11b KO (FF) and WT controls (WT) at D10 and D13. Frequency (Y axis) plotted vs. AUC (X axis), means shown by vertical dashed lines. Full data given in Table S8. *(A)* General developmental progression indicators: Ets1 and Lef1 activities increasing, Bcl11b and Spi1 activities decreasing with development. *(B)* AP-1-associated and Sox5 (TCRγδ-associated) regulons. *(C)* Regulons increasing activity in Bcl11b KO after D10 and/ or concentrated in candidate terminal state. *(D)-(H)* Developmental distribution of regulon activation states, shown in UMAP plots of integrated WT and Bcl11b KO cells, colored with intensities showing relative activities of the indicated regulons. *(D)* Stem/ progenitor regulons (cf. Fig. 4A, Fig. S4A). *(E)* Cell cycle-associated regulon Ybx1 (cf. Figs. S5, S6). *(F)* Developmental progression-associated regulons (cf. panel A). *(G)* AP-1 and Sox5 regulons (cf. panel B). (H) Other Bcl11b KO-induced regulons (cf. panel C). *(I)* Proposed pathway of transcriptional regulatory cascade leading Bcl11b KO cells to alternative developmental endpoint, inferred from Figs. 6, 7 and Fig. S9.

### SCENIC evidence for a cascade of drivers of the Bcl11b KO pathway

Traditional gene network analysis has left unresolved the question of why Bcl11b KO pro-T cells upregulate a set of signature genes that are normally silent, even in pro-T cells before Bcl11b is present to repress them(*34*). Although a known innate-cell regulator, *Id2*, is upregulated even in Bcl11b KO cluster 5 (arrow, Fig. S8D), it encodes a non-DNA binding antagonist and thus cannot directly mediate this novel positive regulation. The cells might be undergoing a mixture of continuing progression (Bcl11b-independent events), failure to complete exit from phase 1 (abnormal progenitor regulator persistence), and also de novo activation of alternative programs (new driver activation), but the regulators involved needed to be identified. We used SCENIC analysis to shed light on other cryptic transcriptional drivers for this process (effects by regulon, Table S8).

Comparison of the WT and Bcl11b KO samples at D10 and D13 showed regulons that exemplified each of these behaviors (Fig. 7A-H). First, *Ets1* and *Lef1* both normally increase expression at commitment, starting immediately after *Bcl11b*(*11, 19, 42*). Their regulons underwent similar increasing activation from D10 to D13 in WT and Bcl11b KO samples alike (Fig. 7A, left). To this extent, both genotypes progressed. Second, the Bcl11a and Spi1 regulons originally operating before commitment lost activity in both, but the decrease appeared faster in WT than in KO samples (Fig. 7A, right). This could reflect weaker silencing of phase 1 regulators in the KO (a few cells might also bypass the normal pathway toward commitment: dashed trajectory in Fig. 6E). Third, multiple regulons showed increased activity in the KO, some starting from the D10 timepoint (Fig. 7D) and others emerging only at D13 (Fig. 7B, C).

Notably, these regulons were largely distinct from the ones that marked developmental progression during phase 1; most appeared newly activated (cf. Fig. 4A, Fig. S5). In Bcl11b KO cells, prominently active by D10 and increasing activity at D13 were regulons associated with Fosb, Jun, and Junb bZIP family members, i.e. components of the classic signal-activated AP-1 factor (Fig. 7B). By contrast, in phase 1, even low-level inferred Jun regulon activity was restricted to Tcf7 KO cells (Fig. 4A, Table S4A). Ikzf2 and Sox5 regulons were also upregulated slightly in Bcl11b KO cells at D10, but very strongly by D13 (Fig. 7B,C). Among normal T lineage cells, Sox5 expression is highly specific for γδ cells, similar to Sox13 (*42*). Between D10 and D13, some Bcl11b KO cells also markedly activated regulons for Nfil3, Rora, and the bZIP factor Maf (Fig. 7C). This genotype-specific upregulation of new regulons suggests a cascade of TFs, possibly signal-activated, that are unleashed in pro-T cells if Bcl11b is absent. The patterns of activity of these regulons projected onto the UMAP1-2 space (Fig 7D-H) provided support for the branched trajectory shown in Fig. 6E.

### Destiny of cells after ‘realizing’ their T-lineage developmental defect

Bcl11b KO pro-T populations abnormally express B-, NK-, other innate-, and some progenitor-associated genes as well as increased γδ-specific genes(*34, 37*), but it has been unclear if these were all expressed in the same cells or segregated among different abnormal subsets. In the UMAP1-2 framework, the regulons upregulated early by Bcl11b KO cells (Jun and Sox5 regulons, Fig. 7G) affected cells throughout the KO-enriched part of the UMAP1-2 space, while activities of the later-activated Ikzf2, Nfil3, Maf and Rora regulons were progressively confined to an extreme ‘tip’ (Fig. 7G, H; cluster 8 in Fig. 5F). Like these regulons, cell type-restricted genes specifically upregulated in Bcl11b KO cells spread across the KO population distribution, but nearly all reached their highest levels at the same tip. Cells most highly expressing NK-(*Il2rb*). ILC-(*Rora, Zbtb16*), and TCRγδ-associated (*Cd163l1*) genes were alike concentrated in this region (Fig. 6B). Thus, the diverse regulators activated in Bcl11b KO appeared to be compatible with convergence on a common regulatory state.

In one respect, this ‘tip of exit’ was qualitatively distinct from the Bcl11b KO intermediate states. Cells in this tip concertedly downregulated both WT-enriched and Bcl11b KO-enriched genes that were Notch-dependent. This was most evident when Bcl11b KO cells only were selected (through sample hashtags) and displayed on Bcl11b-KO-specific UMAPs, with early clusters computationally excluded (Fig S9A-E). Here, proliferating cells are shown on the right (Fig. S9B) while the left margin of the distribution represents late-stage Bcl11b KO cells, expressing *Ikzf2, Cd163l1*, and *Zbtb16* (Fig S9E). In the UMAP3 dimension, cells in the ‘tip of exit’ were distinguished by their downregulation of Notch target genes, not only KO- and WT-shared *Il2ra* and *Ptcra* but also KO-enriched *Nrarp* (Fig. S9C, D). These genes and *Rag1* may be silenced by the maximal expression of *Id2* in this tip, which inhibits bHLH E protein TFs (*46, 47*). Thus, the end-stage of the Bcl11b KO pathway may involve not only the upregulation effects of multiple regulators used in alternative lineages, but also, in one D13 terminus of the Bcl11b KO program (the ‘tip’), the effects of downregulated Notch response (Fig. 7I).

Thus, pro-T cells reaching the threshold of commitment, when they would normally upregulate Bcl11b, responded to the lack of functional Bcl11b by activating an AP-1-like response and progressively upregulating ‘abnormal’ genes in an alternative path to a new regulatory endpoint (Fig. 7I). The cells lacking Bcl11b did not regress along the normal developmental trajectory (Figs. 1B, 4D), but rather spun out into an aberrant maturation program that finally led cells to exit the pro-T pathway via downregulation of Notch signaling responses and further upregulation of innate lineage-associated genes.

## Discussion

Cell fate commitment represents a discrete milestone in a developmental process, an irreversible transition for multipotent cells, resulting in the loss of most of their lineage potentials. However, for a stem cell-derived population that is regulated under an incompletely deterministic gene regulatory network, the process must be viewed as a developmental continuum. In early T cell development there is even marked variation among clones in differentiation speeds leading to commitment(*14*). The process involves multiple TFs (*5, 15, 19, 20, 34, 48–50*), and their individual roles to bring about commitment and to make commitment such a discontinuity have remained poorly defined. Previous studies have studied perturbations of some high-impact regulators either in later stages of development or in slightly different settings, such as fetal liver precursors(*24, 25, 28, 34, 37, 37–40, 51*). To date, however, the lack of single-cell resolution has hindered the full ordering of gene regulatory stages or defining roles of individual TFs in these stages. The stages leading up to commitment have been especially obscure. Previously, our highly sensitive imaging-based seqFISH analysis(*11*) showed that most individual ETP cells in vivo still co-express legacy progenitor genes after activating the critical Notch-induced T cell regulatory genes, *Gata3* and *Tcf7*. Here, we used scRNA-seq coupled with specifically designed and optimized perturbations to dissect the cooperative and antagonistic relationships among key regulators within single early T cell precursors. Specifically, we clarified the roles of 5 TFs leading up to lineage commitment, and identified a separate cascade of poised antagonists restrained by Bcl11b as it directs and enforces lineage commitment.

During the ETP progression to DN2 stage, perturbation of individual TFs clearly shifted differentiation kinetics even without altering the canonical trajectory, and confirmed that the progenitor-associated TFs as well as the “T-lineage” TFs are actively regulating developmental progression speed. Disruption of *Bcl11a*, *Spi1*, or *Erg* accelerated the differentiation process while KOs of *Gata3* or *Tcf7* stalled it. While we failed to resolve effects of Hoxa9 and Meis1 KOs, the Erg KO also shifted the cells to an aberrant, highly proliferative altered-DN2 state marked by high *Klf4* expression and activity, at least in these conditions where apoptosis was blocked. The results imply that such progenitor-associated regulators naturally hold back differentiation progression, while Gata3 and Tcf7 promote differentiation, with Gata3 becoming especially important after the role of Tcf7. This opposition creates an unstable balance between the differentiation-promoting gene network regulators and the stem/progenitor-associated gene network regulators throughout the precommitment stages that determines developmental speed at each step.

SCENIC analysis sharply distinguished effects of these individual TFs, not only as they influence differentiation state but also how they directly or indirectly control alternative developmental pathways, proliferation, Myc activity, and cytoskeleton modules, even when they work in the same direction overall. Importantly, SCENIC also indicated potential indirect effects via expression of other TFs or changes in signaling that alter existing TF activities. Erg was uniquely required to suppress *Klf4* and *Rxra* activity, and restrain excess metabolic and proliferative activation. Bcl11a and Spi1 had similar inferred effects slowing T-lineage progression and transcription of TCR-γδ genes. Notably, however, both also repressed genes associated with NK or ILC lineages, consistent with earlier evidence that Spi1 could inhibit NK fates(*39*). These NK or ILC genes are characteristically repressed by Bcl11b later(*33, 34, 34, 36*), suggesting that they are kept silent by a possible relay of repressive control.

The two required T-lineage factors, Gata3 and Tcf7, differed sharply in their cross-regulatory relationships with Erg and the stem/progenitor factors. Loss of Tcf7 caused uniquely profound inhibition of entry into the T cell program, as found previously(*24, 25, 52*), even in this system supported by anti-apoptotic Bcl2. Despite the shared requirement for Gata3 and Tcf7 in the early T-cell program, the presence of Gata3 tended to suppress proliferation in these conditions while Tcf7 promoted proliferation, an effect seen even though effects of the Gata3-specific sgRNAs in this system were fairly weak. Tcf7 positively regulates *Gata3*, *Il2ra*, and *Bcl11b* as well as multiple T-lineage genes(*24, 52, 53*), and its artificial overexpression of *Tcf7* from an early stage can also promote T cell differentiation(*24*). In contrast, despite their need for *Gata3* (refs (*26–28, 54, 55*)), pro-T cells do not tolerate overexpression of *Gata3*(*56, 57*). The Myc and cell cycle regulatory contrast between these two TFs, shown here, can potentially contribute to an explanation.

The impact of Bcl11b during commitment was qualitatively different: rather than pushing development forward or blocking retrograde development, it appeared to govern a pathway switch. Cells proliferating around the time of commitment appeared primed to fly into either of two divergent developmental trajectories, depending on whether Bcl11b was activated; yet both represented a clear departure from the pre-commitment states of ETP-DN2 stage cells. Progenitor-associated genes were not reactivated in the Bcl11b KO. Instead, many factors that were specifically activated in cells ‘realizing’ the effect of the Bcl11b KO had been relatively inactive throughout the phase 1 stages. While an early increase in *Id2* expression was detected at the first separation of WT and KO transcriptome states, SCENIC analysis also implicated Fos/Jun family regulon activation early in the Bcl11b KO pathway. This raises the possibility that a transcriptional or post-transcriptional bZIP factor activation event provided the actual positive regulatory trigger for the cascade of increasingly abnormal gene expression features proceeding over the next days, as the cells then upregulated Ikzf2, Sox family factors, Nfil3, Zbtb16 and Maf factors and/or their regulons.

Most knockout trajectories apparently converged on a distinct novel endpoint, through which Notch response genes were finally turned off. The maximum upregulation of NK, ILC, and even TCRγδ-associated genes occurred apparently in the same cells, representing the most likely exit path to become the NK-like or innate immune cells observed in previous studies(*36, 37*). The terminal upregulation of *Rora* and *Id2* with loss of Notch input, together with high *Gata3*, recall the gene network circuit recently described for ILC2 differentiation from fetal thymocytes(*58*). This suggests that the pro-T cells entering commitment, in which *Bcl11b* is normally activated, are actually pre-primed for at least two alternative programs: a T cell path dependent on Bcl11b and a ready-for-use alternative, innate-like path.

These single-cell transcriptome analyses thus indicate direct and indirect TF mechanisms, order them, and resolve their additive and stage-specific contributions to pathways for T cell and innate lymphoid development.

## MATERIALS AND METHODS

### Study design

This study dissected the roles of specific TFs in the gene networks leading to and implementing T-cell lineage commitment, by acutely perturbing expression of these TFs in primary pro-T cells and analyzing the impacts on single-cell transcriptomes 3-6 days later.

### Biological materials

Cells were from young adult mice with a C57BL/6 background, from our animal colony. OP9-DLL1 co-cultures and mATO cultures were used for in vitro differentiation. Detailed genotypes and conditions for study of pre-commitment and commitment-stage perturbations, and details of cell culture, retroviral transduction, and flow cytometry, are given in Supplementary Materials and Methods. Retroviral vectors transducing sgRNAs were constructed as described in Supplementary Materials and Methods.

### Single-cell mRNA sequencing

10X Chromium single-cell transcriptome analyses were performed and analyzed as described in detail in Supplementary Materials and Methods.

### Data analyses and availability

An in-house pipeline was developed to identify sgRNAs in individual cells. Seurat 3, Monocle 3, and SCENIC were used for transcriptome analyses. Full description is in Supplementary Materials and Methods. Data are deposited in GEO under accession numbers GSE165835 and GSE183026.

### Statistics

Statistical analyses were conducted using Seurat 3, Monocle 3, SCENIC, R scripts, or GraphPad Prism. Statistical tests and significance levels are reported in respective figure legends.

## ACKNOWLEDGMENTS

We thank Jeff Park and Sisi Chen from the Caltech Single Cell Profiling and Engineering Center for providing support for processing 10X Chromium samples, Rochelle Diamond and members of the Caltech Flow Cytometry and Cell Sorting facility for sorting, Ingrid Soto for mouse care, Maria Quiloan for mouse genotyping and supervision, and Igor Antoshechkin and Vijaya Kumar of the Caltech Jacobs Genomics Facility for bulk RNA sequencing. We also thank Gay Crooks and Amélie Montel-Hagen (UCLA) for sharing the mATO system with us.

## AUTHOR CONTRIBUTIONS

Conceptualization: WZ, EVR

Methodology: WZ, FG

Investigation: WZ, FG, MRW, SJ

Funding acquisition: EVR

Supervision: EVR

Writing – original draft: WZ, EVR

Writing – review & editing: WZ, FG, EVR

WZ designed the project, carried out the experiments, analyzed the data, and wrote the paper. FG wrote the in-house bioinformatic pipeline for perturb-seq and hashtag alignment and assignment, helped with analysis and edited the manuscript. MRW and SJ performed preliminary experiments. EVR supervised research, guided the design of the project, analyzed the data and wrote the paper.

## DATA AND MATERIALS AVAILABILITY

All data, code, and materials used in the analysis will be made available to researchers for reproducing or extending the analysis, upon request to the Corresponding Author, without requiring more than a Simple Letter Agreement or the Uniform Biological Materials Transfer Agreement of the NIH Office of Technology Transfer. All genomic sequencing data have been deposited in Gene Expression Omnibus under accession numbers GSE165835 and GSE183026.

## SUPPLEMENTARY MATERIAL

### SUPPLEMENTARY MATERIALS AND METHODS

#### Animals

C57BL/6J, B6.Cg-Tg(BCL2)25Wehi/J(Bcl2-tg), *Vav1*-iCre mice (B6N.Cg-*Commd10*^Tg(Vav1– icre)A2Kio^/J) and B6.Gt(ROSA)26Sortm1.1(CAG-cas9*,-EGFP)Fezh/J (Cas9), mice were purchased from the Jackson Laboratory. B6.*Bcl11b^yfp/yfp^* reporter mice(*32*) were used for bulk RNAseq analysis, and B6.*Bcl11b^fl/fl^* mice(*34, 37*) were both reported previously. All mice were maintained on the B6 background. For CRISPR/Cas9 experiments, BCL2 transgenic mice and Cas9 mice were crossed to generate B6-Cas9/+; +/Bcl2 heterozygotes for each experiment. For Bcl11b experiments, B6.Bcl11b^fl/+^ *Vav1*-iCre heterozygous mice were bred to obtain *Bcl11b*^+/+^ and *Bcl11b*^fl/fl^ *ROSA26R-YFP* mice with *Vav1*-iCre, as previously described(*37*), which are denoted here as WT and Bcl11b KO. Animals used for these experiments were bred and maintained at the Animal Facilities at California Institute of Technology under conventional Specific Pathogen-Free conditions, and animal protocols were reviewed and approved by the Institute Animal Care and Use Committee of California Institute of Technology (Protocol #1445-18G).

#### Cell lines

To provide a microenvironment that supports T-lineage differentiation *in vitro*, we co-cultivated purified BM cells with the OP9-DLL1 or OP9-DLL4 stromal cell lines(*6*), which were originally obtained from J. C. Zúñiga-Pflücker (Sunnybrook Research Institute, University of Toronto), or with MS5-mDLL1 or MS5-mDLL4 stromal cells(*10*), which were obtained from Dr. Gay Crooks (UCLA) and maintained in our laboratory as described in the original reference(*10*). Details of the differentiation cultures are given below under Method Details.

### CELL PURIFICATION AND CULTURE

#### Primary Cell Purification

For *in vitro (ex-vivo)* differentiation of pro-T cells, bone marrow hematopoietic progenitors were used for input. Bone marrow (BM) was removed from the femurs and tibiae of 10-12 week-old mice. Suspensions of BM cells were prepared and stained for lineage markers using biotin-conjugated lineage antibodies: CD3ɛ (eBioscience, clone 145-2C11), CD19 (eBioscience, clone 1D3), B220 (eBioscience, clone RA3-6B2), NK1.1 (eBioscience, clone PK136), CD11b (eBioscience, clone M1/70). CD11c (eBioscience, clone N418), Gr1 (eBioscience, clone RB6-8C5), and Ter119 (eBioscience, clone TER-119), then incubated with streptavidin-coated magnetic beads (Miltenyi Biotec), and passed through a magnetic column (Miltenyi Biotec), denoted as ‘Lin^-^ BM’.

For all scRNA-seq experiments in this study, the freshly isolated Lin^-^ BM cells were immediately further FACS sorted to purify live (7AAD^negative^), CD45^+^ “LSK” cells (Lin^negative^ Scal^high^ c-Kit^high^), as a more stringently purified source of multipotent hematopoietic stem and progenitor cells. All the BM precursors used (Lin- or LSK) were frozen in liquid nitrogen for storage in freeze-down medium containing 10% DMSO, 40% FCS and 50% OP9 medium (defined below under “BM cell differentiation”) before initiating any differentiation assays.

#### Flow Cytometry and Cell Sorting

Unless otherwise noted, flow cytometry analysis and FACS of all samples were carried out using the procedures outlined. Briefly, cultured cells on tissue culture plates and primary cells from thymus were prepared as single cell suspensions, incubated in 2.4G2 Fc blocking solution, stained with respective surface cell markers as indicated, resuspended in HBH, and filtered through a 40 μm nylon mesh. They were then analyzed using a benchtop MacsQuant flow cytometer (Miltenyi Biotec, Auburn, CA) or sorted with a Sony Synergy 3200 cell sorter (Sony Biotechnology, Inc, San Jose, CA) or with a FACSAria Fusion cell sorter (BD Biosciences). All antibodies used in these experiments are standard, commercially available monoclonal reagents widely established to characterize immune cell populations in the mouse. Acquired flow cytometry data were all analyzed with FlowJo software (Tree Star).

#### BM Cell Differentiation

For standard differentiation of pro-T cells, the hematopoietic progenitors were thawed and either cultured on OP9-DLL1 or OP9-DLL4 monolayers using OP9 medium (α-MEM, 20% FBS, 50 μM β-mercaptoethanol, Pen-Step-Glutamine) supplemented with 10 ng/ml of IL-7 (Pepro Tech Inc) and 10 ng/ml of Flt3L (Pepro Tech Inc); or aggregated to artificial thymic organoids with ATO-mDLL1 or ATO-mDLL4, seated at the air-medium interface on a culture insert (Millipore Sigma) in serum-free ATO medium (DMEM-F12, 2X B27, 30 μM Ascorbic acid, Pen-Step-Glutamine) supplemented with 5 ng/ml of IL-7 (Pepro Tech Inc) and 5 ng/ml of Flt3L (Pepro Tech Inc). If viral delivery of gRNA was required at the start of the culture, thawed BM precursors were initially incubated for 20-24 hours in OP9 medium supplemented with 10 ng/ml of SCF (Pepro Tech Inc), 10 ng/ml of IL-7 (Pepro Tech Inc) and 10 ng/ml of Flt3L (Pepro Tech Inc), without stromal cells, to launch the cells into cycle. This primed them for retroviral transduction before initiating co-culture with the stromal cells, as detailed below in the following section.

#### CRISPR/Cas9-mediated Acute Deletion in Precursor Cell Cultures

To generate input cells, Cas9 mice were first bred to Bcl2-tg mice to generate heterozygotes for both transgenes. LSK or Lin^-^BM cells from these Cas9;Bcl2-tg animals were purified and stored as described above. 20-24 hours after thawing and recovery in cytokines, the cells were transduced with retroviral vectors encoding reporters (CFP) and the indicated guide RNAs (sgRNAs) as detailed below, and then seeded to OP9-DLL1 culture. The methods used to generate the virus supernatant and infecting BM cells were described previously(*38*). For infecting LSK precursors for scRNA-seq (CRISPR), different batches of virus were first tested on primary BM precursors to determine the accurate titers (Fig. S3B), and then delivered to target cells in the actual CRISPR pool perturbation experiments at a precise multiplicity of infection (MOI) of 0.5 or 1. For cell-surface phenotypic assays, cells were analyzed after 2-6 days after culture. For scRNA-seq, retrovirus infected Lin^-^CD45^+^c-Kit^hi^CFP^+^ cells were sorted preparatively on a FACSAria Fusion cell sorter (BD Biosciences).

### CONSTRUCT DESIGN AND CLONING

#### Cloning

The retroviral vector backbone used for sgRNA expression cloning was based on previously published E42-dTet(*38*) with the following modifications: 1) Capture sequence 1 (Cap1) was added to the sgRNA scaffold before the termination signal. 2) One nucleotide ‘G’ was deleted before the sgRNA protospacer insertion site (two *AarI* restriction enzyme cutting sites) to allow compatibility with dual sgRNA vector cloning. The cloning was achieved through high-fidelity PCR (primers as shown below) and Gibson assembly, final cloned product containing the human U6 (hU6) promoter, two *AarI* cutting sites, gRNA backbone with Cap1 sequence and mTurquoise2 fluorescent marker, was as shown in bottom middle in Fig.1D.

For dual sgRNA cloning, a ‘donor’ sequence containing gRNA backbone and mouse U6 (mU6) promoter were obtained from a plasmid modified from Vidigal et al. (*59*). Specifically, the capture sequence 2 (Cap2) were added prior to the termination signal of the sgRNA scaffold backbone; we also found the linker sequence between gRNA backbone and mU6 promoter contained a partial sgRNA backbone sequence that hinders the PCR capability and Gibson assembly accuracy, therefore we cloned to remove the partial repeat sequence in the linker region. The cloning was performed through sequential high-fidelity PCR (primers as listed below) and blunt end ligation.

The pool-based dual gRNA cloning was performed similarly to the protocol described in ref. (*59*) with the modified vector and plasmids above (workflow shown in Fig. 1D), and with minor protocol modifications as follow. 1) the ‘Donor sequence’ containing gRNA scaffold - Cap2 - modified linker – mU6 were obtained through PCR with the modified plasmid, rather than enzymatic digestion. 2) All gel purification steps were avoided, and purifications were achieved with Ampure XP or SPRIselect beads (Beckman Coulter) instead. 3) Selected gRNAs from the oligopool were qPCR quantified before and after the pool-based vector cloning process (Fig. S3A) for quality control, ensuring the amplification evenness of the final plasmid pool. 4) A retroviral vector was used instead of lentiviral vectors.

#### Primers used in cloning

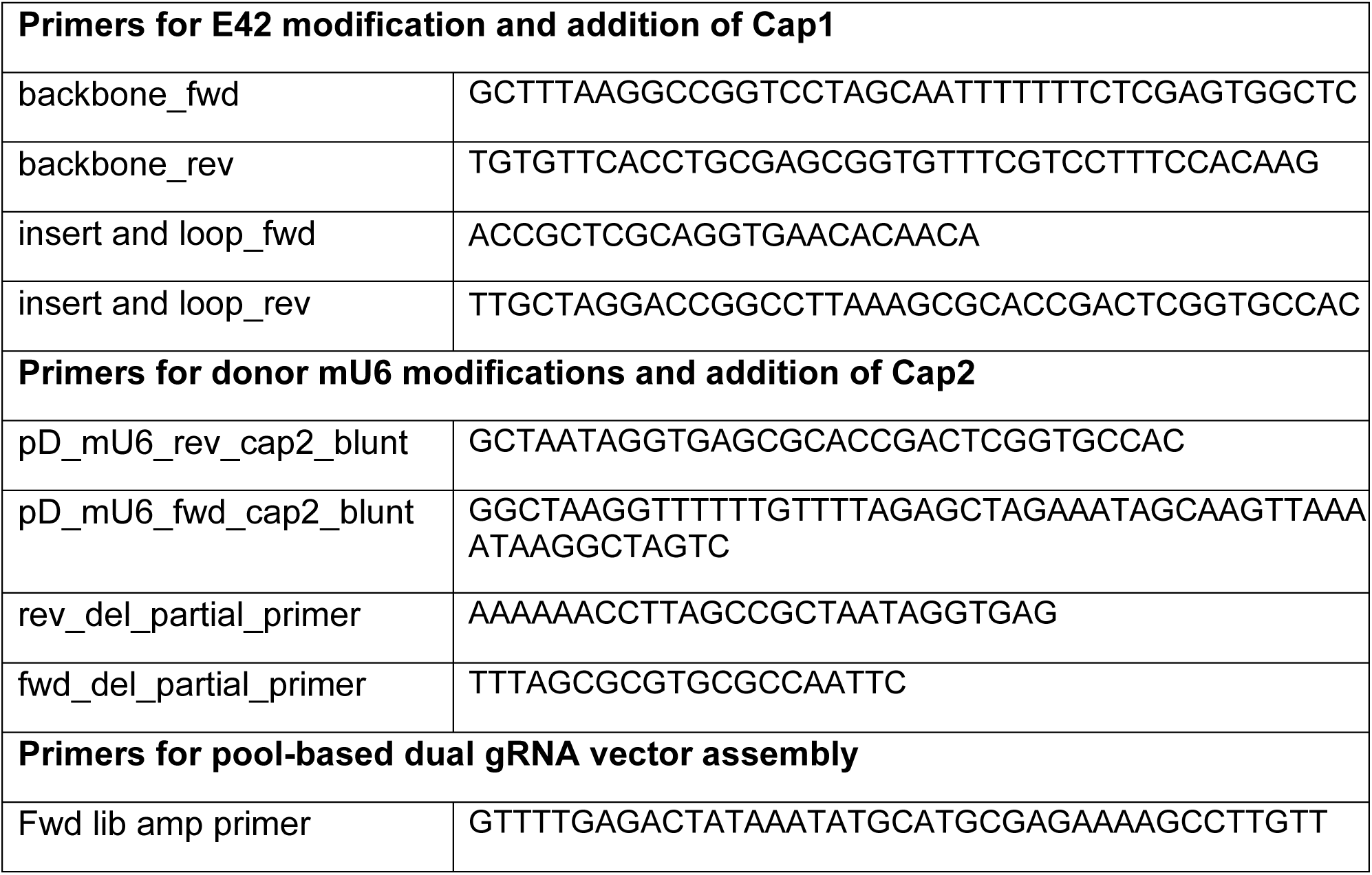
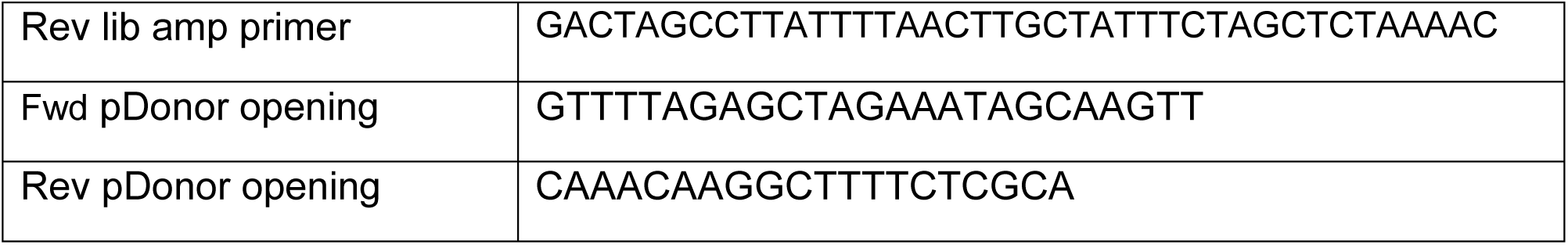

#### sgRNA sequence design and oligo pool design

sgRNAs were designed using GPP sgRNA Designer for CRISPRko (by Broad Institute(*60, 61*)) and GUIDES(*62*) (Sanjana lab) in combination. Specifically, the 3 pairs of sgRNAs of each gene were picked from the top ranked, non-overlapping gRNAs from list generated from GPP designer (https://portals.broadinstitute.org/gpp/public/analysis-tools/sgrna-design). Each pair are designed to target the same exon of the same gene, preferably at or upstream of exons with functional annotation. If GPP designer did not generate all pairs of gRNA sequences desired, GUIDES tool (http://guides.sanjanalab.org/) was used following the instruction. Both designer tools were based on target genome of Mouse GRCm38 and SpyoCas9 (NGG). For oligo pool synthesis, the 110nt oligos were designed and synthesized as follows:

5’—[CATGCGAGAAAAGCCTTGTTTGG]—[gRNA1]—

[GTTTGGGTCTTCGAGAAGACCTCACCG]—[gRNA2]—

[GTTTTAGAGCTAGAAATAGC]—3’

Because the circular DNA will be digested with restriction enzyme *BbsI*, 2 cutting sites of *BbsI* are incorporated in the middle sequence between gRNA1 and gRNA2. Before assembling the sequence of the designed oligo pool, we made sure that the gRNA1, gRNA2 and junction sequences with the left and right primer handles didn’t introduce additional *BbsI* binding sites (‘GAAGAC’, ‘GTCTTC’). For control sequences, we included a pair of non-targeting control sequences from mouse GeCKO library A(*63*), one pair of sgRNA sequences designed against a non-expressing gene Hes3. Note that the final cloning product of the vector library also included a third detectable nontargeting vector that was actually from residual un-cut vector backbone. Cells found to be infected with this construct were included among the ‘controls’ in the differential gene expression analysis but not included in the comparison of differential cell recovery, because their level of representation in the initial pool was unknown. Details of the 110nt designed oligo pool sequences and gRNA sequences are included in Table S9.

### RNA-SEQ AND SINGLE-CELL RNA-SEQ METHODS

#### Bulk RNAseq Analysis for in vivo vs. in vitro reference analysis

For the test of *in vitro* vs. *in vivo* comparability (Fig. S1), Lin^-^ BM cells were harvested from B6.*Bcl11b^yfp/yfp^* animals, and cultured in differentiation conditions as described above. Upon harvesting, cells were subdivided into CD25^low^ for DN1, Bcl11b-YFP^neg^ CD25^hi^ for DN2a, and Bcl11b-YFP^pos^CD25^hi^ DN2a. fractions, followed by RNA purification following the instructions of the RNeasy Micro Kit (Qiagen 74004). cDNA from each sample was prepared using NEBNext Ultra RNA Library Prep Kit for Illumina (E7530, NEB). All bulk libraries were sequenced on Illumina HiSeq2500 in single read mode with the read length of 50 nt. Base calls were performed with RTA 1.13.48.0 followed by conversion to FASTQ with bcl2fastq 1.8.4 and produced approximately 30 million reads per sample.

RNA-seq reads were mapped onto the mouse genome build GRCm38/mm10 using STAR (v2.4.0) and were post-processed with RSEM (v1.2.25; http://deweylab.github.io/RSEM/) according to the settings in the ENCODE long-rna-seq-pipeline (https://github.com/ENCODE-DCC/long-rna-seq-pipeline/blob/master/DAC/STAR_RSEM.sh), with the minor modifications that the setting ‘–output-genome-bam–sampling-for-bam’ was added to rsem-calculate-expression. STAR and RSEM reference libraries were created from genome build GRCm38/mm10 together with the Ensembl gene model file Mus_musculus.GRCm38.gtf. The resulting bam files were used to create HOMER tag directories (makeTagDirectory with –keepAll setting). For analysis of statistical significance among DEGs, the raw gene counts were derived from each tag directory with ‘analyzeRepeats.pl’ with the ‘–noadj-condenseGenes’ options, followed by the ‘getDiffExpression.pl’ command using EdgeR (v3.6.8; http://bioconductor.org/packages/release/bioc/html/edgeR.html). For data visualization, RPKM normalized reads were derived using the ‘analyzeRepeats.pl’ command with the options ‘–count exons –condenseGenes –rpkm’; genes with an average of RPKM ≥1 across samples were kept, and their RPKM values were processed by log transformation. The normalized datasets were then hierarchically clustered with R hclust function based on Euclidean distance and ‘complete’ linkage. The heatmap is visualized with R pheatmap with log_2_ transformed RPKM data (after adding 0.1 to all values).

#### Single Cell RNA-seq comparing *in vitro* with *in vivo* pro-T cells (10X Chromium V2)

Note that only the scRNA-seq data from Fig. S2 was obtained through 10X Chromium V2, the rest of the scRNA-seq data were obtained through the V3 chemistry.

The early T cells derived in ATO-DLL4 from LSK were sorted as shown in Fig. S2A (bottom). The sample was then washed and resuspended to 1 million cells/mL concentration in HBSS supplemented with 10% FBS and 10 mM HEPES, 17,400 cells were loaded into a 10X Chromium v2 lane, and the subsequent preparation was conducted following the instruction manual of 10X Chromium v2. The cDNA library and final library after index preparation were checked with bioanalyzer (High Sensitivity DNA reagents, Agilent Technology #5067-4626; Agilent 2100 Bioanalyzer) for quality control. Following the library preparation, the sequencing was performed with paired-end sequencing of 150nt each end on one lane of HiSeq4000 per sample, by Fulgent Genetics, Inc. (Temple City, CA). The reads were mapped onto the mouse genome Ensembl gene model file Mus_musculus.GRCm38.gtf using a standard CellRanger pipeline. Cells were sequenced to a targeted depth of 50,000 reads per cell.

#### Direct-capture Perturbation scRNA-seq (with 10X Chromium V3)

LSK were purified, recovered in cytokines and infected with MOI 0.5-1, and cultured with OP9-DL1 as described above. Note that two separate packagings of the viral pools were infected into cells at MOI=1.0 or MOI=0.5, in parallel, in separate wells, to serve as biological replicates. The cells infected with independently packaged viruses were cultured separately, in parallel, until harvest, when they could be hashtagged and then finally combined for a pooled scRNA-seq analysis. This strategy was designed to allow capture of a full range of genuine biological response variation among the separate replicates while precluding technical batch effects at the analytical stage.

The medium was changed on day 3. On day 5, the cells were harvested through scraping, filtered and prepared for FACS sorting as described above. Specifically, cells derived from each animal and each time point were stained with a biotin-conjugated lineage cocktail (TCRβ (ebioscience, clone H57-597), TCRγδ (eBioscience, clone GL-3), CD19, NK1.1, CD11b, CD11c, and Gr1). Secondary surface staining was performed with fluorescently conjugated streptavidin, CD45, c-Kit (eBioscience, clone 2B8), CD44 (eBioscience, clone IM7), CD25 (eBioscience or Biolegend, clone PC61.5), and TotalSeq A (Biolegend) anti-Mouse Hashtag 1-6 (1:50, in separate infected samples). A viability dye 7AAD (eBioscience) was again applied to exclude dead cells. The sorted cells (CD45^positive^Lin^low^7AAD^negative^CFP^high^c-Kit^positive^) washed 2 times with HBSS supplemented with 10% FBS and 10 mM HEPES, pooled to target equal cell number from each Hash-tagged sample, and loaded onto one lane of a 10X Chromium V3 chip. The cDNA preparation was performed following the instruction manual of 10X Chromium v3 for perturbation with minor modifications, and the hashtag library was prepared following the Biolegend TotalseqA guide. The cDNA, gRNA library, Hashtag library, and final libraries after index preparation were checked with bioanalyzer (High Sensitivity DNA reagents, Agilent Technology #5067-4626; Agilent 2100 Bioanalyzer) for quality control. All libraries were sequenced on HiSeq4000, by Fulgent Genetics, Inc. Cells were sequenced to at least medium depth of 50,000 reads per cell for cDNA, 20M reads/sample for hashtags and 20M reads/sample for gRNAs.

#### Dual gRNA validation

gRNA2 with cap2 sequence were poorly detected in the pool based perturb-seq setup. We validated the efficiency of gRNA perturbation at position 2 and the dual gRNA perturbation efficacy compared to single gRNA. Specifically, 2 different sgRNA sequences against Il2ra (encoding CD25) were designed, sequences as shown below. Dual gRNA vectors were constructed with combinations of the two gRNAs against CD25 as well as two non-targeting control sequences, as shown in Fig. S3C.

CD25 gRNA1: AGATGAAGTGTGGGAAAACG

CD25 gRNA2: CAAGAGAATCTATCATTTCG

Cont 1: GCGAGGTATTCGGCTCCGCG

Cont 2: GCTTTCACGGAGGTTCGACG

The dual vectors on the same retroviral backbone were packaged individually and delivered to Cas9 expressing Lin^-^ BM precursors for acute knockout, as described above. The cells were cultured for 6 days on OP9-DL1, and analyzed with flow cytometry. Specifically, cells were first stained with a biotin-conjugated lineage cocktail (TCRβ, TCRγδ, CD19, NK1.1, CD11b, CD11c, and Gr1). Secondary surface staining was performed with fluorescently conjugated streptavidin, CD45, c-Kit (eBioscience, clone 2B8), CD44 (eBioscience, clone IM7), CD25 (eBioscience or Biolegend, clone PC61.5), Thy1.2 (Biolegend, clone 53-2.1), and viability dye 7AAD (eBioscience) was again applied to exclude dead cells. The CD25 surface expression were quantified within the 7AAD^-^Lin^-^CD45^+^CFP^+^Thy1.2^+^ population, from each vector, as shown in Fig. S3D. The result confirmed that normally 99% of this population express CD25, and a single gRNA can lead to 70-90% perturbation outcomes. The position 2 on the vector, despite being poorly detected, was surprisingly more effective than position 1 in terms of perturbation efficacy. Importantly, dual gRNAs were consistently more effective in perturbation than the single gRNA vectors. Interestingly, a recent study by Replogle et. al.(*18*) has mentioned the preference of incorporating capture sequence 1 into the gRNA backbone stem loop instead of before the termination sequence to improve the efficiency of perturbation for pool-based CRISPRi scRNA-seq analysis. We have therefore also made the retroviral vector backbone with cap1 moved to the stem loop region, and the perturbation efficiency is pending future validation.

#### Single Cell RNA-seq (10X Chromium V3) for comparison of Bcl11b KO and WT Samples with Cell Hashing

LSK from Bcl11b WT and KO animals were obtained, aliquoted into 6-7k cell/tube, and stored in liquid nitrogen as described above (individual animals were not pooled). To setup the culture, cells were thawed and aggregated with MS5-mDLL4 (800-1000 LSK and 150k MS5-DLL4 cells per ATO), and seeded on culture inserts as described above. The ATO medium was changed every 3-4 days. After culturing for 10-13 days (note experiment 1 had only D10, and experiment 2 had both D10 and D13), the ATO was mechanically disrupted and *ex-vivo* derived T cells were prepared for FACS sorting as described above. Specifically, cells derived from each animal and each time point were stained with a biotin-conjugated lineage cocktail (TCRγδ (eBioscience, clone GL-3), CD19, NK1.1, CD11b, CD11c, and Gr1). Secondary surface staining was performed with fluorescently conjugated streptavidin, CD45, c-Kit (eBioscience, clone 2B8), CD44 (eBioscience, clone IM7), CD25 (eBioscience or Biolegend, clone PC61.5), and TotalSeq A (Biolegend) anti-Mouse Hashtag 1-8 (1:50, in separate samples). A viability dye 7AAD (eBioscience) was applied to exclude dead cells. The sorted cells (CD45^positive^Lin^low^7AAD^negative^CD25^positive^) washed 2 times with HBSS supplemented with 10% FBS and 10 mM HEPES, pooled to target equal cell number from each Hash-tagged sample, and loaded onto one lane of a 10X Chromium V3 chip. The cDNA preparation was performed following the instruction manual of 10X Chromium v3, and the hashtag library was prepared following the Biolegend TotalseqA guide. The cDNA, tag library, and final library after index preparation were checked with bioanalyzer (High Sensitivity DNA reagents, Agilent Technology #5067-4626; Agilent 2100 Bioanalyzer) for quality control. The cDNA final libraries was sequenced on HiSeq4000 or NovaSeq 6000, and the tag library was sequenced on HiSeq4000, by Fulgent Genetics, Inc. Cells were sequenced to an average depth of 50,000-70,000 reads per cell for cDNA and ∼2,500 reads per cell for hashtags.

### DATA ANALYSIS

#### Mapping of scRNA-seq Sequences, Hashtag and gRNA Identification

Single-cell RNA-seq data were processed using 10X Cellranger 3.0.0 software. Standard cellranger-mm10-3.0.0 reference annotations were loaded to the pipeline for read mapping and gene quantification.

To process single-cell hashtag and guide RNA sequencing data, two ultrafast in-house tools (**hashtag_tool** and **guiderna_tool**) (https://github.com/gaofan83/single_cell_perturb_seq/) were developed to process raw fastq data and generate count tables (Fig.S3E). The results are typically delivered within one minute. Downstream R codes can be used to binarize the count tables for identity assignment using Gaussian Mixed Modeling.

As note, **guiderna_tool** was specifically developed for our dual-guide system with two guide-RNA sites (targeting different sites of the same gene) are engineered in the same vector backbone. Based on 10X bead chemistry, **Capture1** (5’-GCTTTAAGGCCGGTCCTAGCAA −3’) and **Capture2** (5’-GCTCACCTATTAGCGGCTAAGG −3’) sequences recognize expressed **Guide1** and **Guide2** RNA molecules that have reverse complement capture sequences inserted. Specifically, **Capture1** and **Capture2** sequences should pair with **Guide1** and **Guide2**, respectively. From in-house single-cell guideRNA data, UMI counts can be calculated for **Guide1** list of barcodes and **Guide2** list of barcodes. As note, **guiderna_tool** uses both capture sequences in R1 reads and template switching oligo sequence (TSO) in R2 read for read filtering and sorting; then potential **protospacer** sequences in R2 reads (after 5’ TSO sequence) are mapped against the corresponding guide library (**Guide1** or **Guide2**) for quantification. In contrast, **Cellranger** finds a constant region after **protospacer** region in R2 first, then **protospacer** abundances in R2 are calculated. Since **guiderna_tool** utilizes both R1 and R2 read information for filtering, it is expected to be more accurate.

### Gene and Cell Filtering, Data Alignment, and Clustering Analysis

The 10X Chromium V2 scRNA-seq analysis used to compare in vitro vs. in vivo-derived early pro-T cell samples (Fig. S2) was based on data filtered on cells with at least 1200 genes expressed (transcript count over 1); outliers with more than 4300 genes or 23k unique transcripts were removed (potential doublet) from the ATO scRNA-seq dataset, and outliers with more than 4400 genes or 27k unique transcripts were also removed from the thymocyte dataset (10X V2 run1 from ref.(*11*)), and only genes that were found expressed in at least 3 cells were kept in the analysis. The cells were further filtered to keep only cells with mitochondrial RNA content of less than 7.5-9%. These QC filters resulted in 6167 cells in the ATO scRNA-seq sample and 4783 cells in the thymocyte sample in Fig. S2. The top 3000 variable features were identified from each of the two datasets and integrated with CCA algorithm using the 3000 anchor features and 20 dimensions in Seurat v3(*64*). The principal component analysis was performed on the integrated dataset. In Fig.S2B, the UMAP1-2 display was analyzed based on PCs 1-20 of integrated data, except as other. In Fig.S2C-F, UMAP and clustering analyses were performed on integrated data after cell cycle regression according to Seurat 3 instruction. Louvain clustering was performed on the first 20 PCs with the resolution set to be 0.7, and the top 10 enriched genes in each cluster were calculated with Wilcoxon Rank Sum test, shown in the heatmap (Fig.S2F).

For all the 10X Chromium V3 scRNA-seq datasets (all datasets in the study except in Fig.S2), analysis was based on data filtered on cells with at least 1300 genes expressed (transcript count > 1). The doublet elimination was guided through Cell Hashing. Specifically, the number of features vs. number of unique transcripts detected were plotted, and cells with more than 1 Cell Hashing tags were considered doublets and highlighted on the scatterplot. We dropped both the ‘cell hashing identified doublets’ and outliers with only one hashtag identified but which fell in the region of high numbers of gene detected and transcripts detected similarly to ‘cell hashing doublets’. The subsequent integration and clustering analysis were performed similarly described above, using Seurat 3. For Fig. 2, after UMI count matrices for all the samples were merged, scRNA-seq analysis was performed for cells with low mitochondrial content. Normalized UMI counts were further scaled with mitochondrial content and cell cycle stages regressed out using Seurat 3. (ref(*65*)).

Unless specified, the trajectory and pseudo-time analysis with Monocle 3 were all performed on the cells that passed the filtering steps described above.

#### Kullback-Leibler (KL) Divergence Calculation

Kullback-Leibler (KL) divergence was used to compare two probability distributions. Here, we used it to calculate the pair-wise distance of samples based on cluster distributions of each sample. The number of cells in each cluster was counted in each sample, and the counts were aggregated and converted into a cluster distribution matrix. The pair-wise KL divergence matrix were computed with the KL function in philentropy (v 0.4.0) in R using the cluster distribution matrix, and the returned distance matrix is visualized in heatmap format with heatmap.2 using ggplots (v 3.1.0).

#### Pool-based CRISPR/Cas9 KO Assignment Strategy

The guide 1 and guide 2 identified using the in-house pipeline described above, were analyzed for agreement, as shown in Fig. S3F. The result shows general agreement between the paired guides designed, but guide2 UMI counts captured by capture 2 sequence were significantly lower than guide1-capture1, and were often undetected in individual cells. We validated that the guide2-capture2 were very efficient in generating KO phenotype (described above in Methods and in Fig. S3C-D), and Sanger sequencing of sampled plasmids confirmed good sequence quality of the guide1 and guide2 pairing (data not shown). Therefore, to maximize yield of cell numbers, in Fig. 1-4, we used guide1 assignment for gene perturbation identification. Specifically, the binarized gRNA assignment matrix was generated as described above, for guide1-capture1. The cells with 0 guide 1 assignment were represented as ‘unknown’, and 2 assignments or above were identified as ‘multi’ (for potential multiple vector infection). The 3 gRNA pairs (identified through 3 singly infected guide RNA vectors) against the same genes were examined separately for analysis in Fig. 2F and S4B, and aggregated for gene specific KO effects in the rest of the analysis. The cells identified as ‘unknown’ or ‘multi’ (even with multiple infection against the same gene) were dropped from the analysis. Note that we noticed that Erg.3 vector produced slightly different perturbation outcome than the other 2 Erg vectors, therefore only cells with Erg.1 and Erg.2 gRNAs were selected and used to represent Erg KO in differential gene expression and SCENIC analysis.

#### Differential Gene Expression analyses

Genes differentially expressed between different sample classes within a population were defined (Table S3, Table S7) using Seurat 3 with standard criteria of qval < 1E-02, minimum fraction of cells expressing the gene ≥ 0.05, and natural log Fold-change (lnFC) between populations (using that pseudocount) ≥ 0.1. Note, in the analysis we used the default Seurat 3 pseudocount setting of 1, which reduces noise from low-level transcripts but compresses the dynamic range for comparisons between genes non-expressed and modestly expressed in two different populations. Given the robustness in identifying highly expressed genes’ differential expression pattern, the criteria described above were used for the DEG identification in the paper and for analyses derived from their relative expression levels (e.g. Fig. 3C,D, Fig. 4C, Fig. S4F-I, Fig. S6, Fig. S8). Despite pseudocount of 1 being common practice, the choice of pseudocount is arbitrary and can introduce subtle bias in DEG analysis.

For some future analyses, however, greater sensitivity to differences in expression of lower-level transcripts could be desirable. Although setting pseudocounts close to zero arbitrarily increase the variance of genes with zero counts, we found that the fold change might better reflect lowly expressed transcription factors. Hence, we also provide a separate Table of differentially expressed genes calculated from the pool-perturbation CRISPR experiment using alternative criteria, with qval still <1E-02 and minimum fraction still ≥ 0.05, but with pseudocount set to 0.01 and minimum ln FC ≥ 0.2 (Table S10).

#### SCENIC Analysis and Visualization Graphics

We performed SCENIC(*41*) analysis on the scRNA-seq data without separating individual samples, following the workflow using the default parameters in SCENIC R setup. The co-expression network was generated using GENIE3(*66*), and potential direct-binding targets (regulons) were based on DNA-motif analysis. AUC, which identifies and scores gene regulatory networks or regulons in single cells, was calculated using AUCell as previously desribed(*41*). The motif bindings were inferred based on publicly available motif binding databases provided by the S. Aerts lab. Following the regulon calculation, the individual cells’ regulon AUC values from 3.4_regulonAUC.Rds were extracted and separated by sample identity of perturbations based on gRNA assignment (CRISPR) or hashtags (Bcl11b). The AUC distributions of different genotypes were compared and analyzed for changes in regulon activity between different perturbations. Note that this SCENIC analysis was completely independent of the definition of DEGs by Seurat 3, above. The full summary list of AUC values by samples for each KO are included in Tables S4 (CRISPR pool perturbation) and S8 (Bcl11b experiment 2).

#### Software Details

The bulk RNA-seq analysis were mapped with STAR (v2.4.0) and post-processed with RSEM (v1.2.25). The Fastq outputs for scRNA-seq data were aligned with Cellranger 3.0.0. The scRNA-seq downstream analysis were performed mainly in R (version 4.0.2) with the following packages: ggplot2(v3.3.2), dplyr(v1.0.2), cowplot(1.1.0), Seurat(v3.2.2), AUCell(v1.10.0), RcisTarget(v1.8.0), GENIE3(v1.10.0), SCENIC(v1.2.2), monocle3(v0.2.3.0), ggraph(v2.0.4), igraph(v1.2.6), philentropy (v 0.4.0), gplots (v 3.1.0). The statistical analyses were performed with methods as specified in the text using software mentioned above or with Graphpad Prism (v9.0.0).

## Supplementary Figures and Legends

**Fig. S1.**
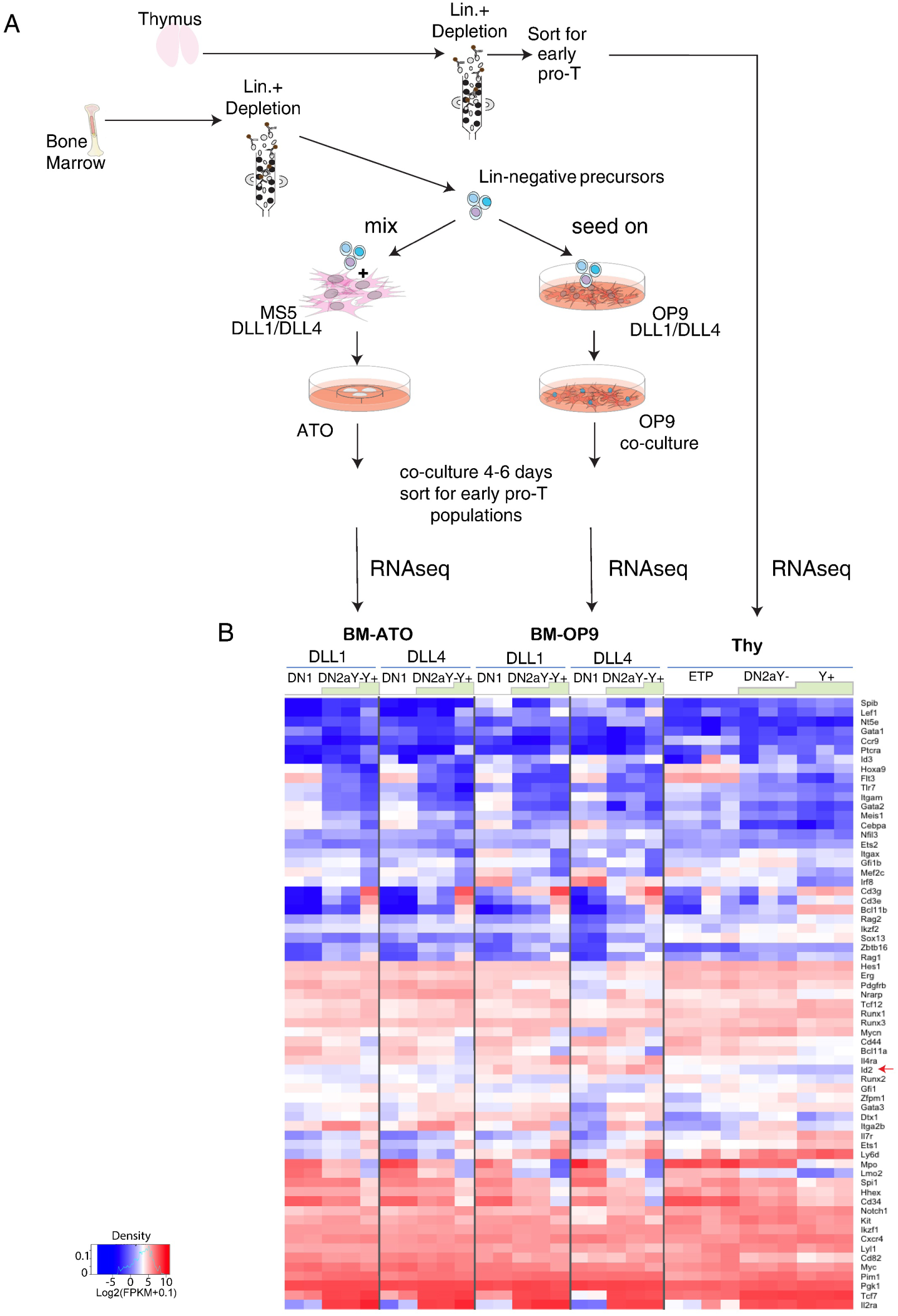
Bulk RNA-seq analysis of *in vivo* and *ex-vivo* derived early pro-T cells. *A)* Illustration of early pro-T cells harvested from thymus or derived by *in vitro* differentiation from bone marrow (BM) precursors. The *in vitro* culture systems shown include both OP9-DL co-culture systems with OP9-DLL1 or OP9-DLL4 stromal cells, and 3D artificial thymic organoid (ATO) systems based on co-aggregation with MS5-mDLL1 or MS5-mDLL4 stromal cells, as detailed in Methods. *B)* Expression heatmap of bulk RNA-seq measurements, comparing corresponding subsets of early pro-T cells harvested *in vivo* with those sorted from *in vitro* differentiation of BM precursors in the culture systems as illustrated in A. All genes plotted are from a curated list of developmentally important regulatory genes described in a previous study(*11*). Color scales indicate raw expression levels as log_2_(RPKM+0.1), without row normalization. This provides a direct visualization of the full dynamic range of gene expression levels.

**Fig. S2.**
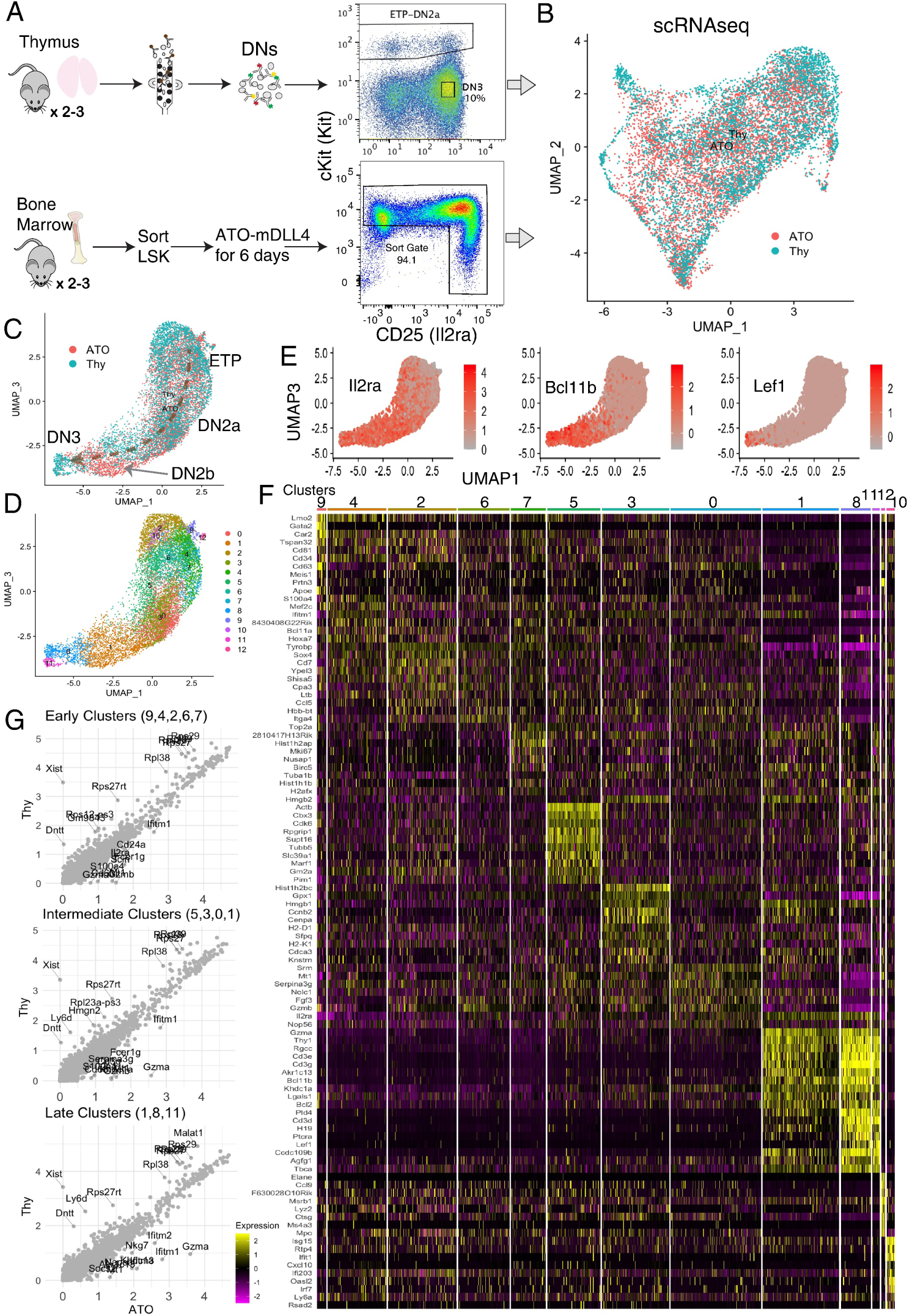
Gene expression profiles of BM-derived early pro-T cells in vitro recapitulate those of *in vivo* thymic early pro-T cells, on single-cell level. *(A)* Illustration of sample purification procedures and FACS sorting strategies for the scRNA-seq experiments, comparing *in vivo* and *in vitro* derived early pro-T cells’ single-cell expression profiles. *(B)* Aligned *in vivo* and *in vitro* derived scRNA-seq profiles after CCA scaling, shown in UMAP1-2. The integrated data contains 6167 cells from ATO, and 4783 cells from thymus, 10950 cells total. *(C-F)* Analysis of cell cycle-regressed integrated data, shown in UMAP 1-3 for the clearest separation of developmental stages. *(C)* Cells colored by origin of sample, i.e. ‘Thy’ for *in vivo* thymocytes and ‘ATO’ for pro-T cells derived from ATO-DLL4 as discussed in A). Trajectory annotated based on expression of stage marker genes. *(D)* Cells colored with clustering assignment of integrated data. Clustering was performed with Louvain clustering algorithm on PC1-20 of integrated data. *(E)* T-developmental marker genes expression pattern. *(F)* Heatmap displaying the top 10 enriched genes in each sub-cluster ordered by approximate developmental progression based on gene expression and connectivity in low dimensional displays. (Seurat 3 pipeline with minimum fraction of expressing cells ≥ 0.25, pseudocount 1, Wilcoxon rank sum test with avg_logFC threshold of 0.3). *G)* Within the early, middle and late developmental sub-clusters, the average gene expression level (size and log normalized transcript count data) comparison between thymic pro-T cells (Thy) and the *in vitro* derived pro-T cells (ATO). Off-diagonal outlier genes were labeled. Note that the ‘Thy’ data were obtained from female mice whereas the ‘ATO’ data were derived from LSKs of male mice, hence the Xist expression in ‘Thy’ data only.

**Fig. S3.**
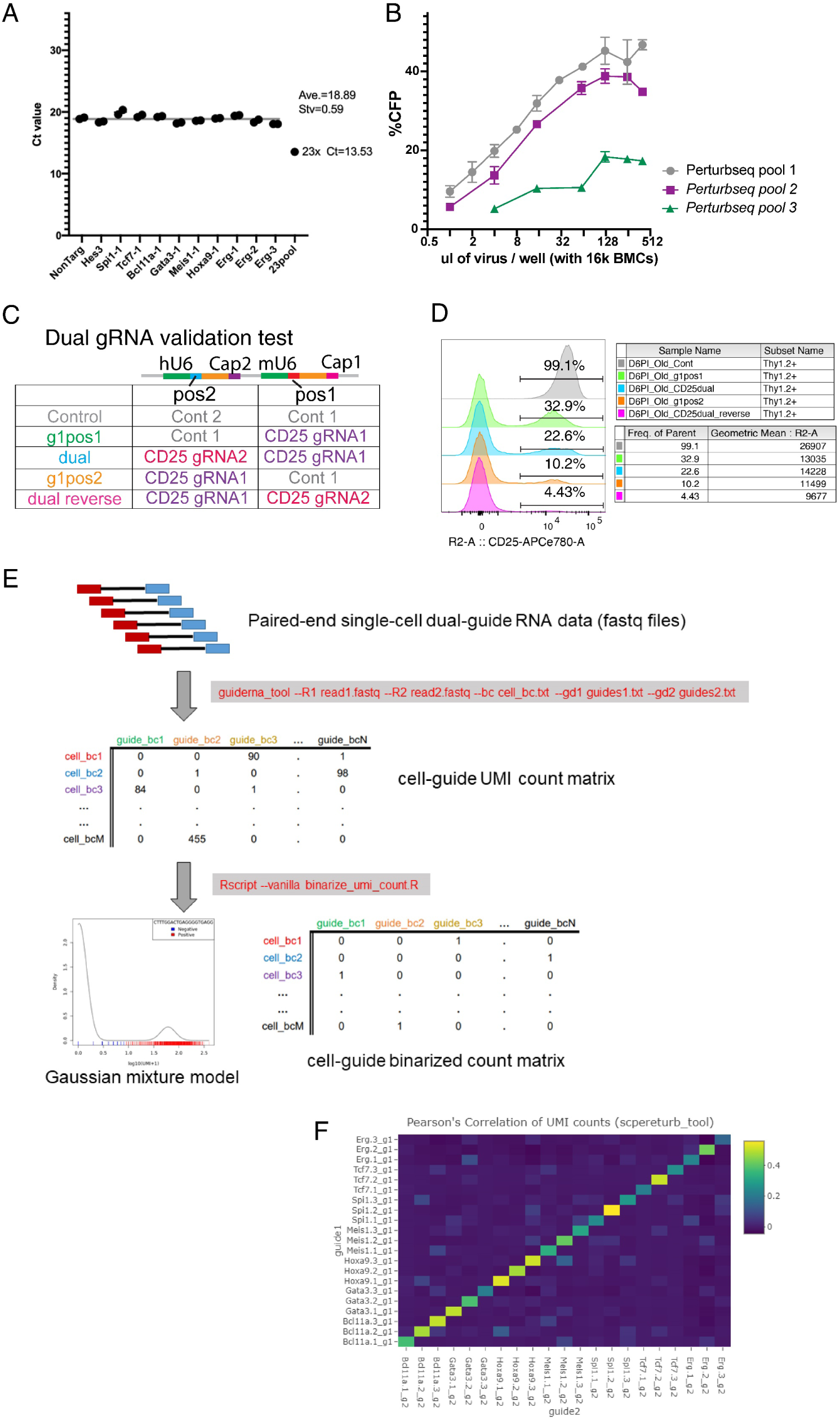
Technical and analytical details of dual gRNA perturb-seq. (*A)* The qPCR C_T_ values of selected vectors from plasmid pool, which sample the evenness of plasmid vector backbones in the cloned dual-gRNA pool. The result showed an evenly synthesized pool of plasmids that was used for viral packaging. (*B)* Multiplicity of infection (MOI) titration with the viral pools containing dual gRNA vectors. All the batches of viral titers were tested separately on primary BM cells, to target precisely MOI of 0.5-1 for the scRNA-seq experiment (viral usage ∼ 40-64% of the plateau). Because of the inferior infectivity of pool 3, only pool 1 and pool 2 were used in this study. *(C)-(D)* The validation experiment of dual gRNA effectiveness of acute gene perturbation. *(C)* In response to the low recovery of guide2-cap2 read counts (from pos2 on the illustration), tested directly whether the gRNAs from pos2 were adding effectiveness to the acute perturbation in our system. Here, *Il2ra* (encoding CD25) was targeted by two gRNAs with switching positions. *(D)* Flow cytometry analysis of the dual gRNA validation test. Results shown indicate that the dual gRNA perturbation efficiency towards the same gene was consistently better than a single gRNA. In fact, the same gRNA sequence in pos2 was actually even more effective in perturbation compared to pos1. *(E)* In-house bioinformatic processing pipeline to align the dual gRNA with both Cap1 and Cap2 information (detailed in Methods). *F)* Pearson correlation of UMI counts from guide1 and guide2 assignment.

**Fig. S4.**
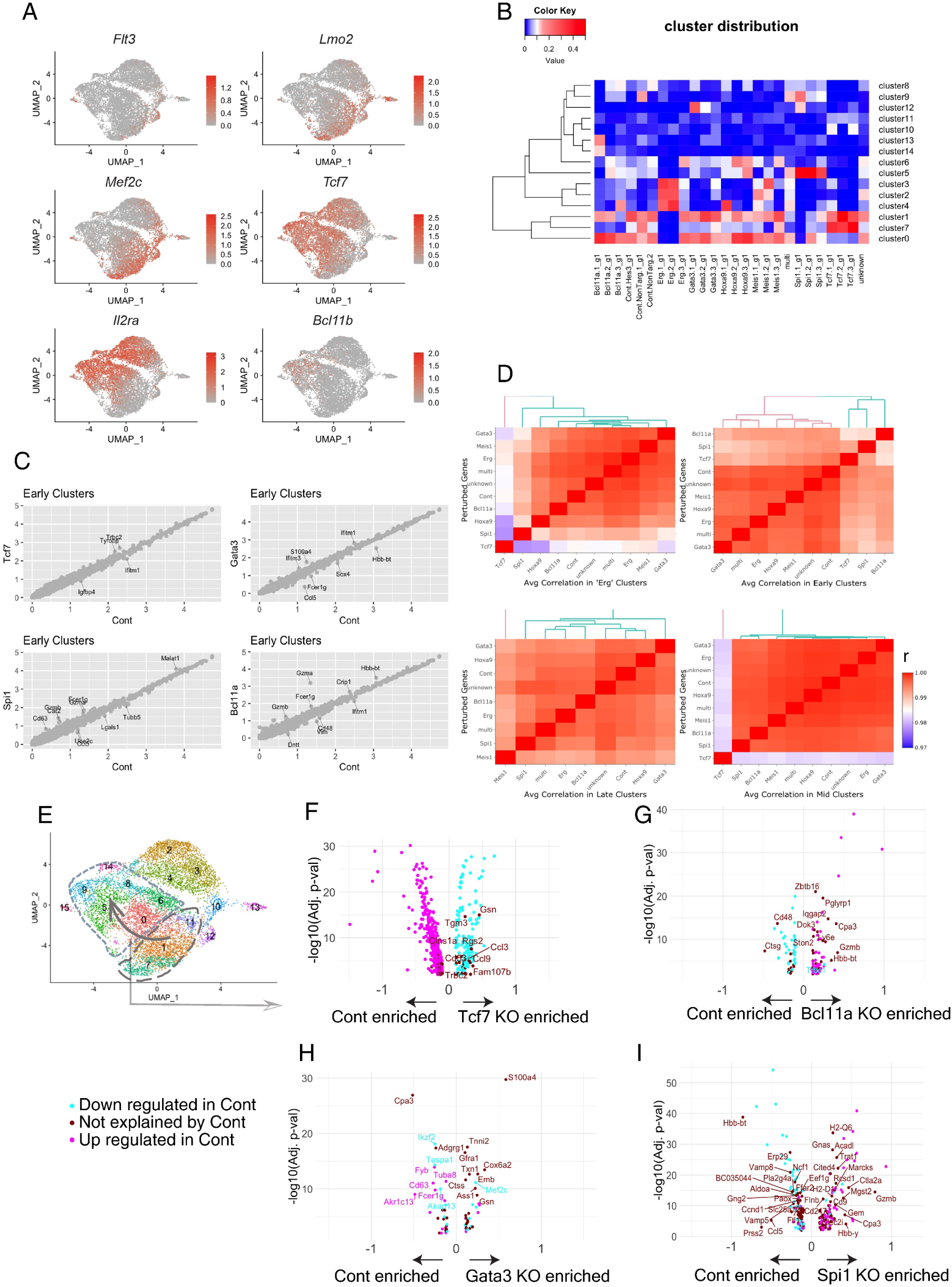
Relationship of knockout effects to the normal T-cell developmental trajectory. *(A)* UMAP1-2 plots of whole pool-perturbation ensemble with cells highlighted based on expression of major T-cell developmental landmark genes. *Flt3, Lmo2, Mef2c*: progenitor-associated genes. *Tcf7*: onset at earliest T-cell specification. *Il2ra*: onset is ETP to DN2a transition marker. *Bcl11b*: activated only during T-lineage commitment, in DN2a to DN2b transition. *B)* Low dimensional representations and clustering assignments mostly represent the differences between genotypes (KOs). Heatmap of cluster distributions of individual dual gRNA perturbations. Color scale shows fractions of cells in each cluster. Result show a general agreement of cluster distributions of perturbations against the same genes, with a few exceptions, such as Erg.3 (3^rd^ pair of gRNAs targeting Erg). *(C)* Scatterplots comparing average gene expression levels [ln(count+1)] between KOs and Control (Cont), in the shared early common clusters (7,1,11 in sub-clusters defined in Fig. 2A). The results show high similarity of gene expression levels between KOs and Cont in the same ‘common clusters’. values in ln(count+1) *(D)* Heatmaps showing Pearson correlations between average gene expression patterns in different perturbations, when they fall in the shared ‘common clusters’. (The 4 ‘common clusters’ were defined as: ‘early clusters’: 7, 1, 11; ‘mid clusters’: 0, 6; ‘late clusters’: 9, 5, 8; ‘Erg clusters’: 4, 3, 2 sub-clusters defined in Fig. 2A, respectively.) Note that the color scale of heatmap represents Pearson correlation of 0.97 to 1. This implies within the same ‘common clusters’, the expression profiles between KOs and Cont are very similar. *(E)-(I)* Dissection of KO effects on developmental “speed” (cf. Fig. 3B): relationship of genes differentially regulated in each KO to genes normally changing expression in development *(E)* UMAP1-2 illustration of clusters pooled and contrasted for the differential expression test performed on Cont cells only, to reveal genes that change during the early to late transition under normal developmental condition (i.e., no perturbation). (*F-I)* Volcano plots showing the differentially expressed genes between Cont and individual KOs (x axes), with the color of dots representing whether the gene normally increased or decreased expression in Cont cells only during the normal developmental progression from ‘early clusters’ to ‘mid’ and ‘late’ clusters, as annotated in E. Specifically, genes that were upregulated during normal early to late transition in Cont cells are labeled in cyan, and the downregulated genes are labeled in magenta. Genes that were differentially expressed between KOs and Cont but not differentially regulated in this normal developmental transition are labeled as dark brick red. Concordance of color labels with positive or negative values along the x axis help to visualize whether the differential gene expression between WT and KOs was merely reflecting a developmental acceleration or a stalled progression.

**Fig. S5.**
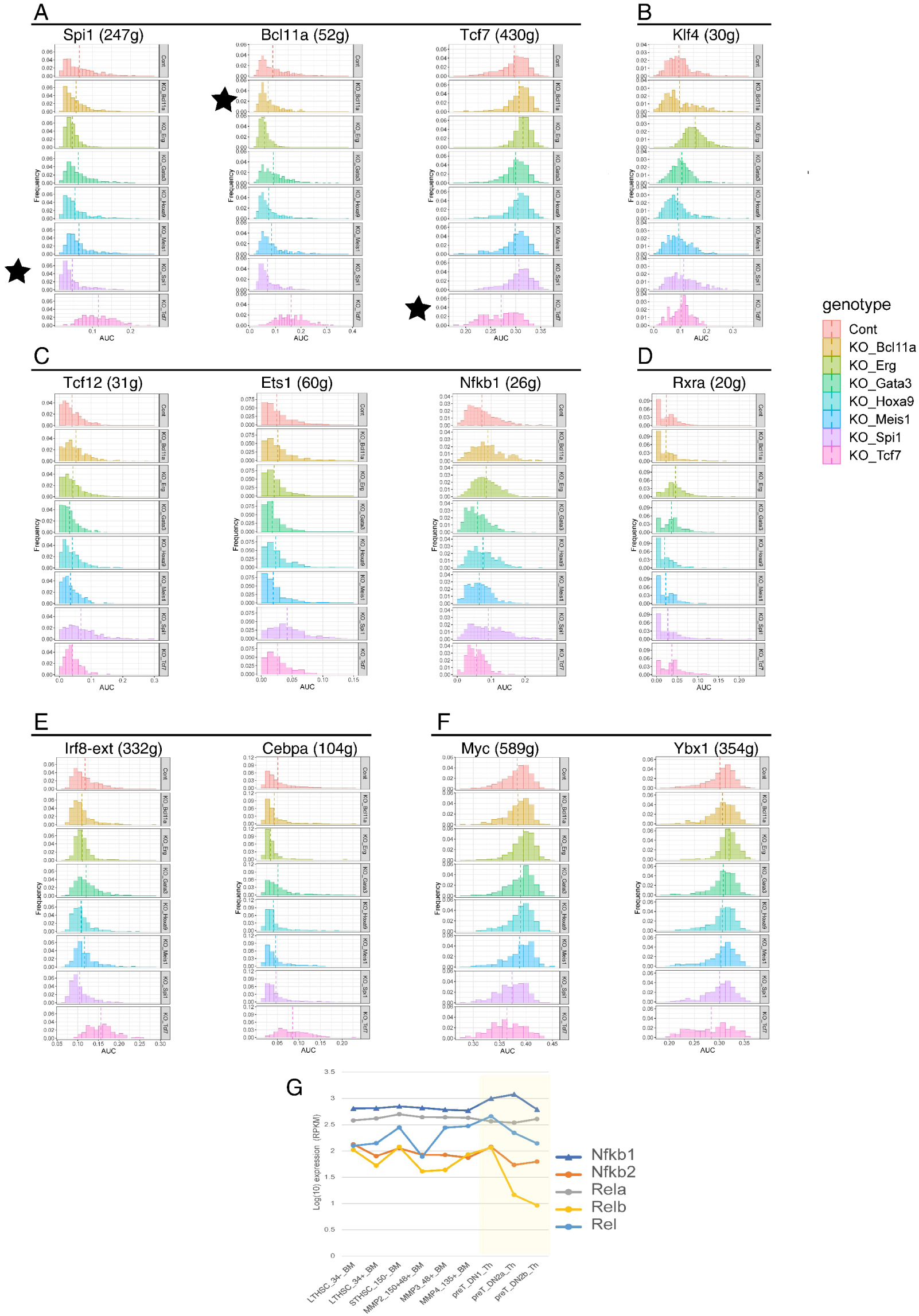
Inferred TF activities and regulatory connections by SCENIC: histograms of individual representative regulons in the 8 CRISPR pool-perturbation genotypes. SCENIC analysis was performed on subsets of Cont and individual KOs. *(A)-(F)* Histograms of frequencies of cells with a given Area Under the Curve (AUC) of activity of the indicated regulon, among cells of a given genotype (perturbation state) within the pool-perturbation scRNA-seq ensemble. Vertical dashed lines indicate means. *(A)* Regulons corresponding to factors directly targeted among the perturbation conditions, indicated by black stars. *(B)* Klf4 regulon, specifically upregulated when Erg is removed. *(C)* Regulons specifically upregulated in normal cells progressing along the T-cell developmental pathway. Note that all are specifically upregulated when Spi1 is removed. *(D)* Rxra regulon, rarely expressed but specifically upregulated in Erg KO cells. *(E)* Myeloid-associated regulons for Irf family transcription factors and Cebpa: Note strong upregulation in absence of Tcf7 and possibly in absence of Gata3 as well. *(F)* Cell blastogenesis and proliferation-associated regulons, for Myc and Ybx1. Note upshift in Erg KO and downshift in Tcf7 KO. Analysis details are given in Methods. For full quantitation of all scorable regulons in the dataset, see Table S4. *(G)* Strong increase in Nfkb1 regulon activity from ETP to late DN2a/DN2b pro-T cells (Fig. 4A) contrasts with near-maximal expression in ETP stage of genes encoding NF-κB components themselves. Shown are RNA-seq expression values of the five major NF-κB subunit-encoding genes from hematopoietic stem cells to DN2b cells, from data of the Immunological Genome Project consortium(*42*).

**Fig. S6.**
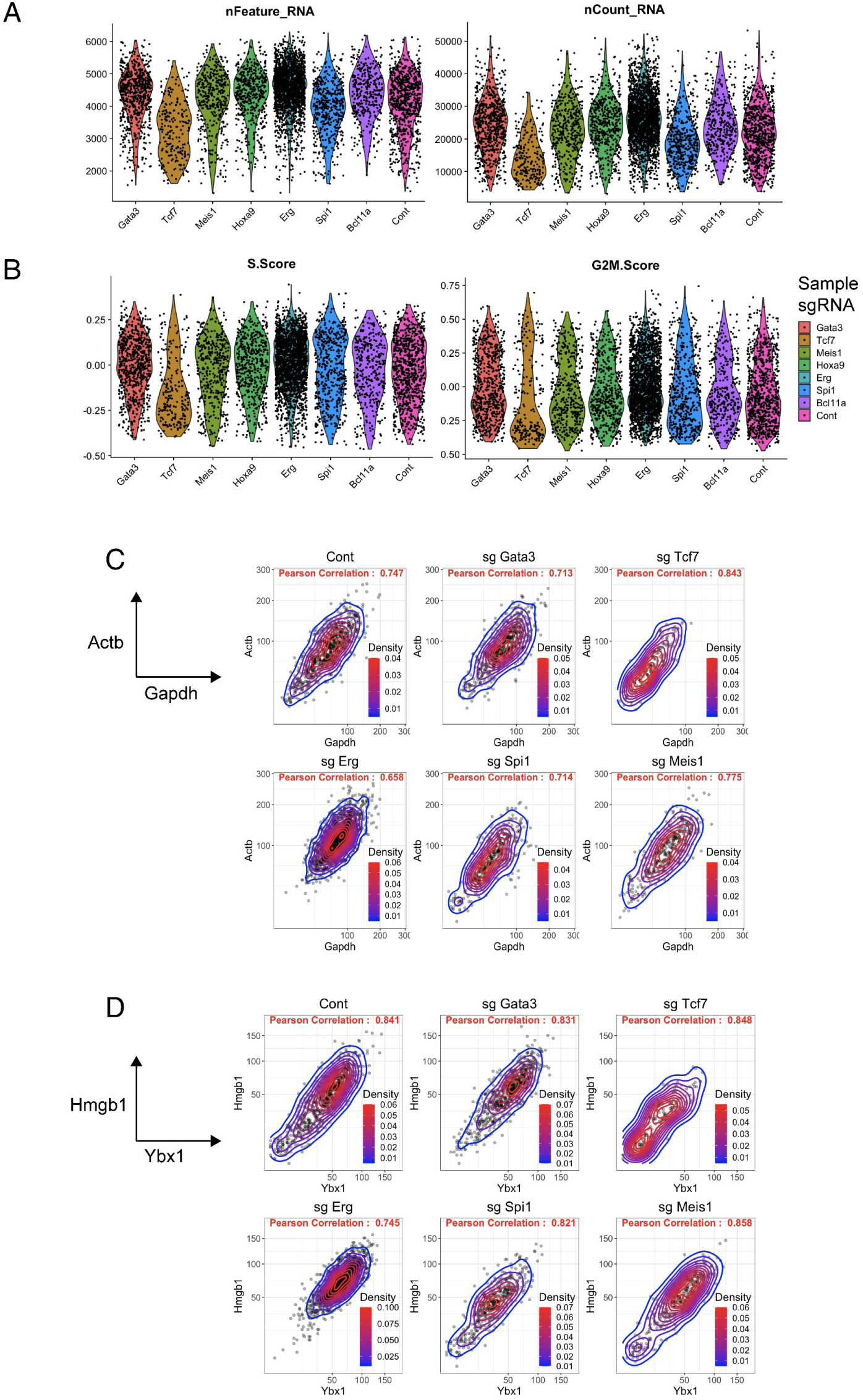
Detailed evidence of TF regulated activities in developing pro-T cells. *(A)* Distributions of number of genes detected, number of transcripts detected, and *(B)* inferred cell cycle stages, in subsets of the pool-perturbation ensemble separated according to genes perturbed. *(C)-(D)* Scatterplots of transcript distributions of housekeeping genes *(C)* and cell cycle-associated genes *(D)*, separated by genes perturbed. Number of cells of each KO in analysis: Cont: 793, Bcl11a: 353, Erg: 1397, Gata3: 686, Hoxa9: 503, Meis1: 409, Spi1: 481, Tcf7: 215.

**Fig. S7.**
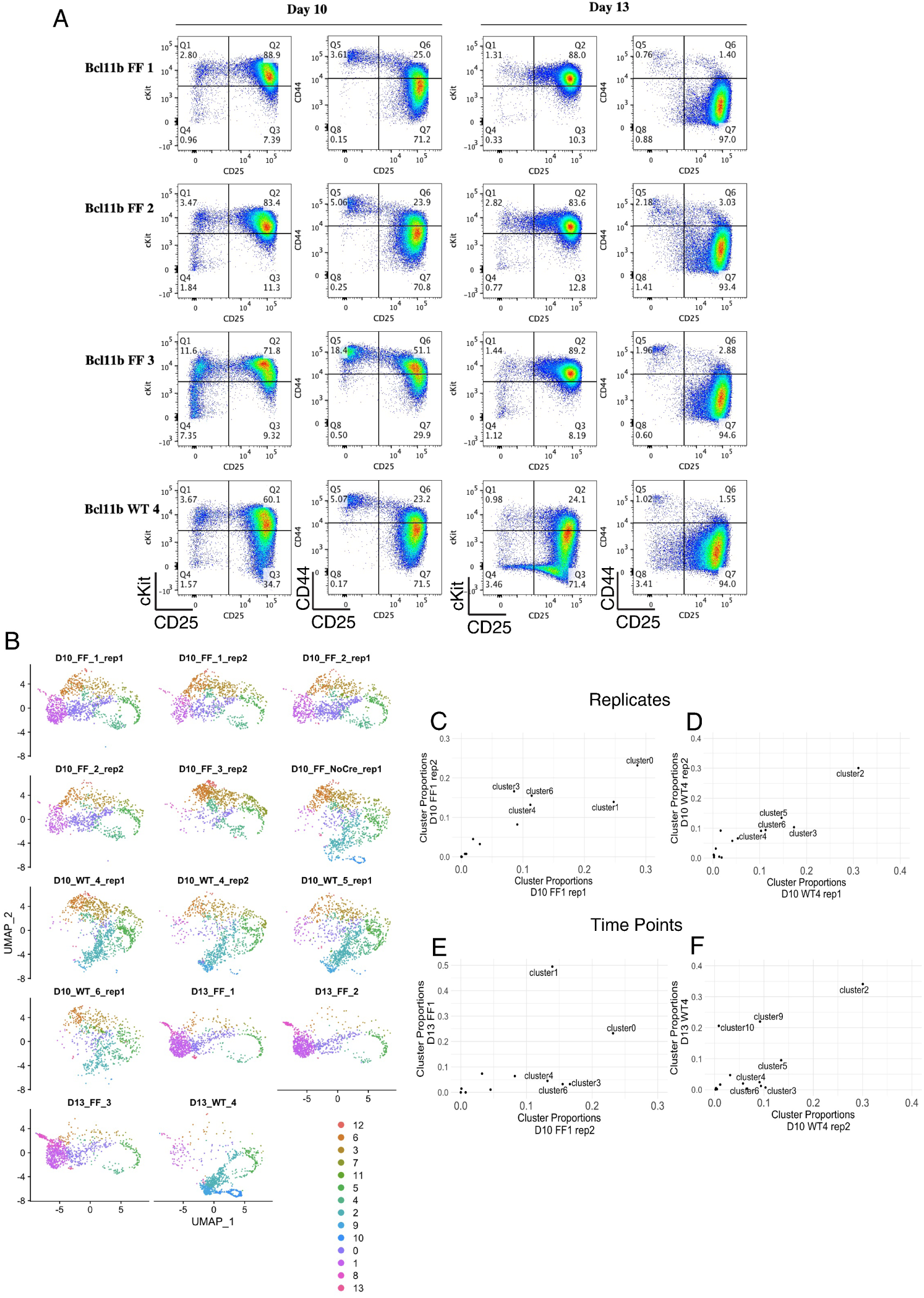
Surface marker and transcriptome profiles of WT and Bcl11b KO single-cell samples. (*A)* Flow cytometry profiles of WT and Bcl11b KO cells (labeled as ‘FF’ for Bcl11b homozygous flx/flx locus) collected from ATO-DLL4 culture system at D10 and D13 of culture. All cells contain *Vav1-iCre*, so the Bcl11b locus designation determines whether the cells will be functionally WT or KO. Note that the *Bcl11b* locus is starting to be transcribed in normal DN2A cells around D7. CD25 staining from the indicated samples is shown correlated with Kit (c-Kit in figure) and CD44 staining. Data show that compared to WT control, the cells missing Bcl11b could still similarly turn off CD44 but failed to downregulate c-Kit expression levels. Cell numbers in subpanels: ‘D10_FF_1_rep1’: 1366, ‘D10_FF_1_rep2’: 1083, ‘D10_FF_2_rep1’: 1261, ‘D10_FF_2_rep2’: 1038, ‘D10_FF_3_rep2’: 1115, ‘D10_FF_NoCre_rep1’: 1300, ‘D10_WT_4_rep1’: 1352, ‘D10_WT_4_rep2’: 1075, ‘D10_WT_5_rep1’: 1377, ‘D10_WT_6_rep1’: 795, ‘D13_FF_1’: 1017, ‘D13_FF_2’ : 1061, ‘D13_FF_3’: 1022, D13_WT_4 : 1147. *(B)* UMAP 1-2 display of the integrated scRNA-seq data, separated by individual samples. *(C)-(D)* Cluster distributions (shown in proportions), comparing different biological replicates of cells derived from BM of the same animal origin. This shows replicability of *ex-vivo* derivation and scRNA-seq experimental setups. R=0.89 for C, R=0.94 for D. *(E)-(F)* Cluster distributions comparing samples harvested from D13 (y axis) with those harvested from D10 (x axis) of the same animal origins. WT: R=0.66, FF: R=0.54. Cluster assignment is the same as described in Fig. 5F.

**Fig. S8.**
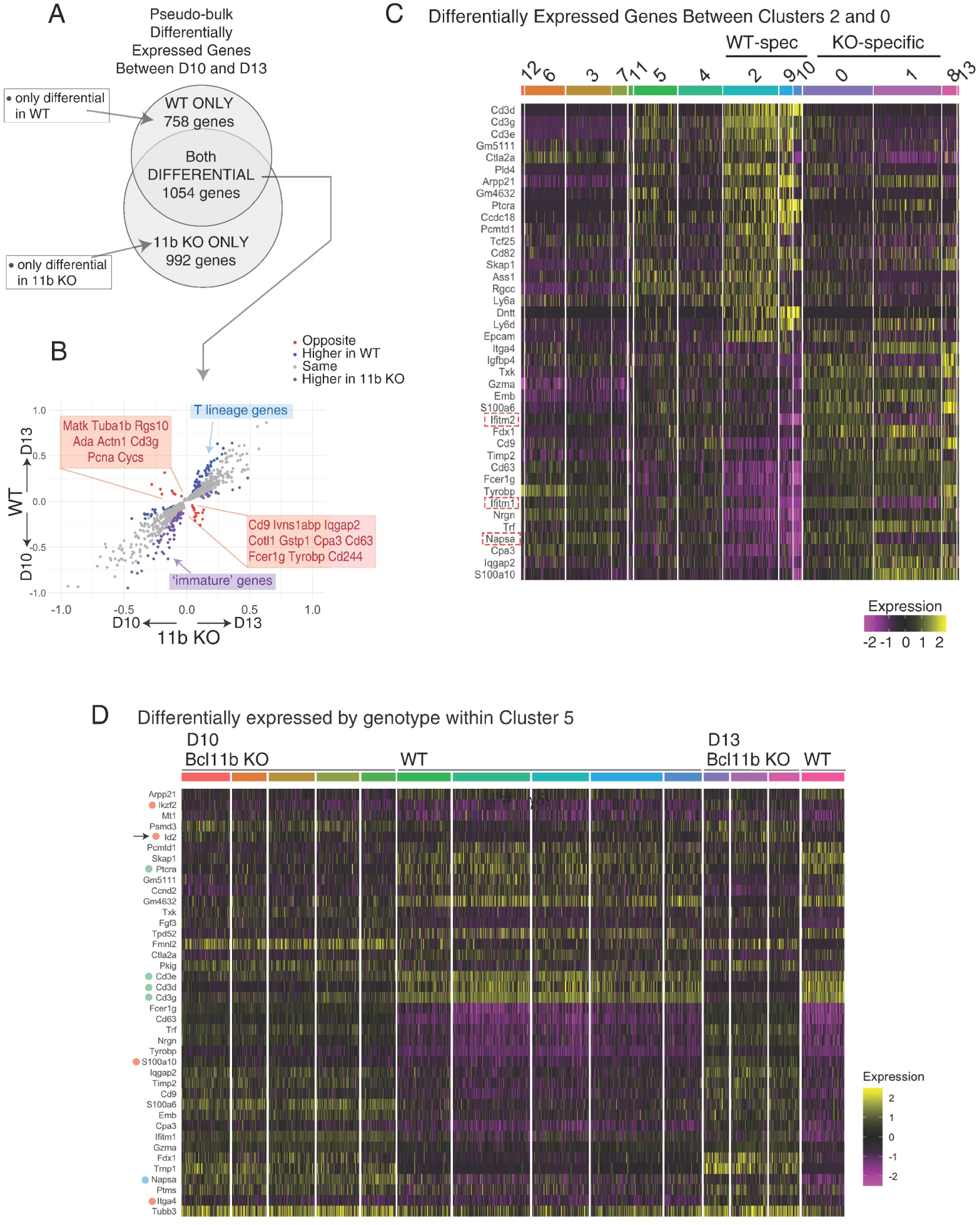
Time-dependent gene expression deviations between Bcl11b KO and WT late pro-T cells. *(A)* Venn diagram showing the numbers of genes with changing expression from D10 to D13 and their overlaps between the two genotypes. These genes are listed in Table S6. Differential expression comparison was made between genes in cells that were harvested on D10 and D13, separately analyzed in WT and Bcl11b KO, and the Venn diagram shows the intersection between these two DEG sets. Differential expression tests were performed with a generalized linear model with Quasi-Poisson distribution of transcript count using Monocle3. Statistical significance was calculated by Wald test, and adjusted p-values represent false discovery rate. The differentially expressed genes were defined based on adj.p-val < 1E-10 for Bcl11b KO and adj.p-val < 1E-5 for WT, accounting for the cell number difference. WT: N= 5899 cells from D10 and 1147 cells from D13. Bcl11b KO: N= 5863 cells from D10 and 3100 cells from D13. *(B)* Scatterplot comparing genes that were differentially regulated in both the WT and Bcl11b KO pseudo-bulk measurements, showing whether the directions of expression change over time were the same or opposite in the two genotypes. Shown are the 1054 genes that were significantly differentially expressed in both of the genotypes. Axes represent “log fold changes” of Seurat-processed expression with respect to time in each genotype. Red dots: genes that changed expression in opposite directions between WT and Bcl11b KO; cyan dots: genes expressed more highly in WT at both timepoints (≥1.7 fold difference in ‘estimates’, and the ‘estimate’ in at least one of the genotype ≥0.1); blue-purple dots: genes with sustained higher expression in Bcl11b KO (≥1.7 fold difference in ‘estimates’, and the ‘estimate’ in at least one of the genotype ≥ 0.1). *(C)* Heatmap showing the top 20 differentially expressed genes between cluster 0 and cluster 2, in both directions, also see Table S7A. (Wilcoxon Rank Sum test, filtered by minimally expressed by 25% cells in of one of the clusters, and adjusted p-val < 1e-50). *(D)* Heatmap of differentially expressed genes between WT and Bcl11b KO in only the cells from cluster 5, revealing the gene expression differences that caused the separation shown in Fig. 6F. (Wilcoxon Rank Sum test, filtered by minimally expressed by 25% cells in of one of the clusters, and adjusted p-val < 1E-20, top and bottom 20 genes ranked by average log expression differences (‘avg_logFC’ in Seurat) are displayed, calculated using Seurat 3. Full DE analysis see Table S7B.) Red dots: genes that are enriched in Bcl11b KO compared to WT within cluster 5 only, more markedly expressed in D13 than D10. Green dots: T-lineage progression genes, higher in WT. N=762 cells from Bcl11b KO in cluster 5 and N=868 cells from WT in cluster 5.

**Fig. S9.**
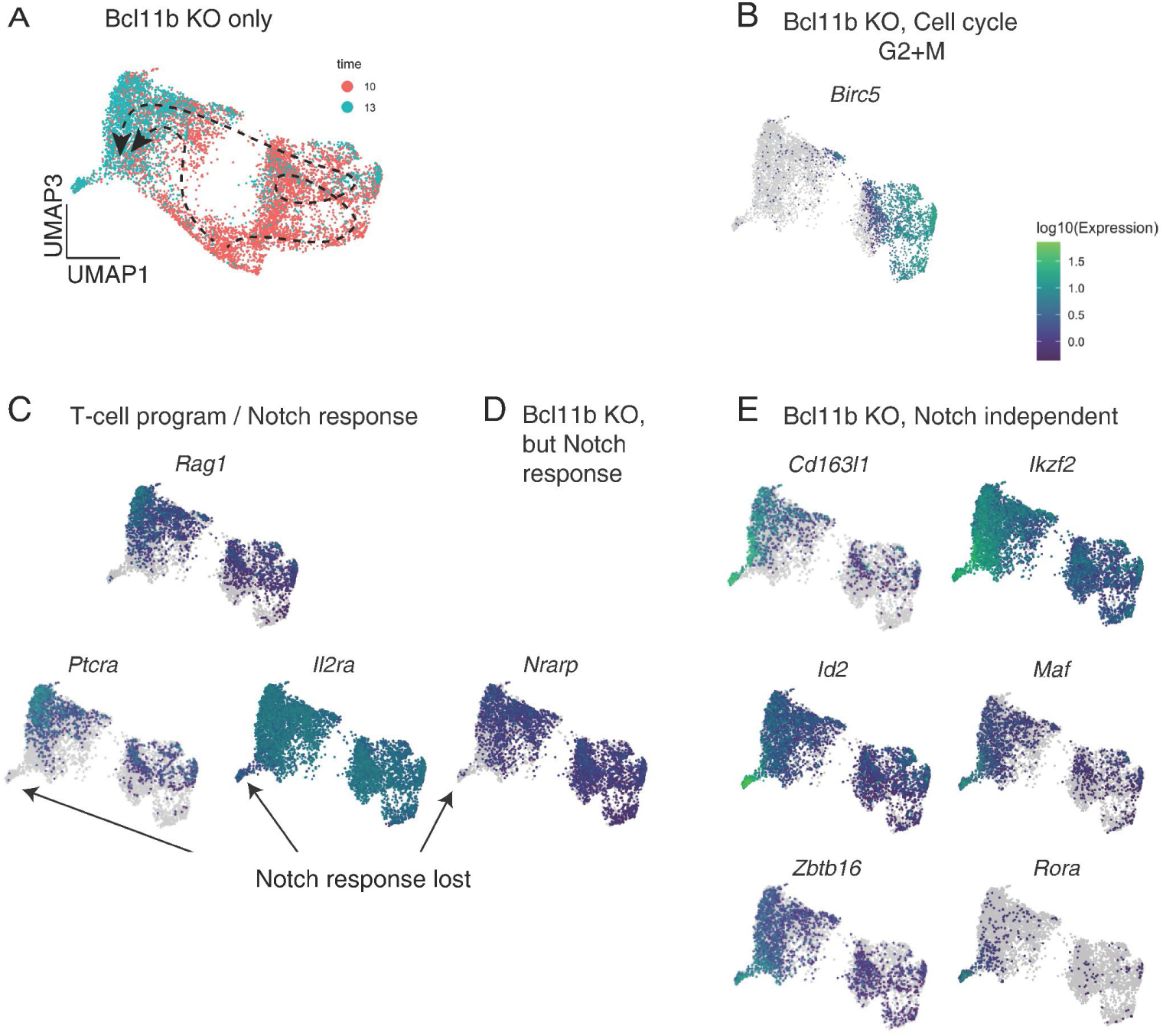
Fine-grained examination of the Bcl11b KO trajectory: identifying accumulation of abnormal gene expression features and a potential exiting point from T-lineage program. *(A)* UMAP 1 vs. UMAP 3 plot of only the cells derived from Bcl11b KO animals from both experiments, D10 (red) and D13 (blue) timepoints. Bcl11b KO cells were first subsetted from the ensemble (cf. Fig. 6B) by the hashtag assignments, excluding the shared non-T cluster that is also present in Bcl11b KO and WT, and then used to generate these UMAP plots (N=8899 cells). *(A)* Cells colored by time point of sample collection, showing Bcl11b KO samples’ distribution changes with respect to time, on UMAP1-3. Central gap in pattern: cells with Bcl11b KO genotype from D10-specific clusters representing stages before the normal time of Bcl11b expression were included to compute UMAP layout, but are masked out from these displays to focus on changes in transcriptome distributions at later stages. Dashed lines indicate approximate trajectories for the Bcl11b KO cells (paths shown including more or less proliferation). *(B)-(E)* Selected genes’ expression patterns on the UMAP 1-3 plot of Bcl11b KO cells only. N=6820 cells. *(B)* Expression of *Birc5* as marker for cycling G2+M cells. *(C)* The expression of Notch response genes within the post-commitment T-cell program, *Ptcra* and *Il2ra*, and E protein-regulated *Rag1*: all are downregulated at the bottom left ‘tip’ of this UMAP 1-3 display. *(D) Nrarp*, a Notch-dependent gene that is actually upregulated in Bcl11b KO populations overall, also loses expression in cells at the bottom left ‘tip’. *(E)* Notch-independent genes associated with innate lymphoid cells, TCRγδ cells, and agonist-selected TCRαβ T cells undergoing intrathymic stimulation: all show increased expression in D13 cells (left-hand half of UMAP1-3 plot) with maximal expression at bottom left ‘tip’.

## Supplementary Tables

Tables themselves are submitted as individual datasets.

Table S1. Bulk RNA-seq Differential Expression analysis comparing corresponding subsets of thymocytes and BM derived pro-T cells, generated under OP9-DLL1 and mATO in vitro differentiation conditions.

Table S2. Genes enriched in each sub-cluster of early pro-T pool-perturbation scRNA-seq. Calculations in this Table are based on Seurat 3 with minimum fraction of expressing cells ≥ 0.25, using Wilcoxon rank sum test with avg_logFC threshold of 0.3 (default pseudocount=1).

Table S3. Differential Expression analysis comparing Control (Cont) and seven TF KOs in early pro-T cell pool-perturbation scRNA-seq. Differentially expressed genes in each comparison of KO to Control were defined with qval < 1E-02, minimum Ln Fold Change ≥ 0.1, minimum fraction of cells expressing ≥ 0.05. Standard for Seurat 3, a pseudocount value of 1 was used for undetected transcripts. These calculations provided the input for Fig. 2B and Fig. 3. DEG lists calculated using alternative criteria giving greater sensitivity to differences in low-level expressed genes are presented separately, in Table S10.

Table S4. (A) SCENIC output of effects from early pro-T cell pool-perturbation scRNA-seq on the complete set of regulons scorable in this dataset. (B) Regulon target gene lists in this dataset. Note that SCENIC analysis is based on calculations from the primary data that are completely independent of the Differential Expression analyses.

Table S5. Characterization of gene expression subclusters in WT and Bcl11b KO samples. (A) Marker genes for all subclusters in Integrated WT and Bcl11b KO scRNA-seq analysis; adjusted pval <1E-02. (B) Proportion represented by each subcluster in each of the D10 and D13 WT and Bcl11b KO samples.

Table S6. Differential gene expression analysis between D10 and D13 timepoints in WT and Bcl11b KO, pooled from single-cell data to give “pseudo-bulk” comparisons. DEG criteria as in Table S3.

Table S7. Differentially expressed genes between specific Clusters in WT and Bcl11b KO scRNA-seq analysis. (A) Differentially expressed genes between Cluster 0 and Cluster 2 in integrated WT and Bcl11b KO scRNA-seq analysis. (B) Genotype-based differential expression analysis (WT vs. Bcl11b KO) within shared Cluster 5. DEG criteria as in Table S3.

Table S8. (A) SCENIC analysis of regulon effects of Bcl11b KO as compared to WT at D10 and D13 of culture. (B) Regulon target gene lists from this dataset.

Table S9. Designed gRNA list (A) and pool oligo list (B) used for generating pool-perturbation dual gRNA vectors.

Table S10. Differential Expression analysis in early pro-T cell pool-perturbation scRNA-seq using an alternative method to increase sensitivity to low levels of expression. To increase accuracy of fold change determinations, calculations in this Table are based on use of a lower pseudocount floor value for undetected transcripts (pseudocount=0.01), with qval < 1E-02 and fraction of cells expressing ≥ 0.05. To offset the contribution of low-expression noise, the minimum ln Fold Change was increased to ≥ 0.2.

